# Modeling Mechanical Feedback Mechanisms in a Multiscale Sliding Filament Model of Lymphatic Muscle Pumping

**DOI:** 10.1101/2022.11.30.518078

**Authors:** Peter Y. Xie, Christopher J. Morris, Christopher Bertram, David Zaweija, James E. Moore

## Abstract

The lymphatic system maintains bodily fluid balance by returning interstitial fluid to the venous system. Flow can occur through a combination of extrinsic pumping, due to forces from surrounding tissues, and intrinsic pumping involving contractions of muscle in the lymphatic vessel walls. Lymph transport is important not only for fluid homeostasis, but also for immune function, as lymph is a carrier for immune cells. Lymphatic muscle cells exhibit cardiac-like phasic contractions to generate flow and smooth-muscle-like tonic contractions to regulate flow. Lymphatic vessels therefore act as both active pumps and conduits. Lymphatic vessels are sensitive to mechanical stimuli, including flow-induced shear stresses and pressure-induced vessel stretch. These forces modulate biochemical pathways, leading to changes in intracellular calcium and interaction with regulatory and contractile proteins. In a multiscale computational model of phasic and tonic contractions in lymphatic muscle coupled to a lumped-parameter model of lymphatic pumping, we tested different models of the mechanical feedback mechanisms exhibited by lymphatics in experiments. Models were validated using flow and pressure experiments not used in the models’ construction. The final model shows that with flow-induced shear stress modulation, there is a small change in flow rate but an increase in muscle efficiency. A better understanding of the mechanobiology of lymphatic contractions can help guide future lymphatic vessel experiments, providing a basis for developing better treatments for lymphatic dysfunction.

## 1. Introduction

The lymphatic system is responsible for maintaining fluid balance by returning interstitial fluid to the venous system. Lymph transport occurs due to a combination of extrinsic pumping (external squeezing of lymph vessels) and active contractions of muscle in the lymphatic vessel wall. Interstitial fluid first enters the initial lymphatics, which are bounded by overlapping endothelial cells acting as one-way valves, and do not actively contract. The initial lymphatics converge into larger collecting lymphatic vessels which actively contract due to lymphatic muscle cells (LMCs) lining the vessel wall [1]. Unlike the blood circulatory system, the lymphatic network does not have a central pump. Instead, lymphatics consist of pumps in series – lymphangions (short segments of vessel between two valves), which propel fluid centrally. Lymph transport usually works against an adverse pressure gradient, propelling fluid from low or sub-atmospheric pressures in the interstitium [2,3]. Lymph flow is essential for fluid regulation and adaptive immunity and is implicated in most cancer metastases. Deficiencies in lymph transport can result in lymphedema, an accumulation of fluid and proteins in the interstitium. There is currently no effective cure for lymphedema, and the absence of treatments is partly due to lack of understanding of LMC contraction dynamics [4,5]. Lymphatics are also the route for the advection of antigens, cytokines, and immune cells to lymph nodes for the initiation of the adaptive immune response [6,7]; thus, lack of lymph transport can lead to an inability to deal with infection.

LMCs possess functional and molecular characteristics of both myocardium (cardiac muscle) and blood vessels (smooth muscle) [8,9]. They display both rapid cardiac-like phasic contractions to generate flow and smooth-muscle-like slow, long-lasting tonic contractions (producing constriction to regulate flow). The excitation of LMCs results from the rhythmic spontaneous transient depolarization (STDs) arising at pacemaker sites [10–12]. These depolarizations can sum spatially and temporally to generate action potentials which then cause rapid increases in the intracellular calcium concentration through the opening of voltage-dependent Ca^2+^ channels [13,14]. In both smooth and striated muscle types, changes in intracellular calcium lead to the activation of the muscle regulatory proteins that produce the actin-myosin interactions to contract the muscle cell. Contraction is essentially uniform within a lymphangion (not peristaltic) and is usually coordinated with surrounding lymphangions [15].

### 1.1 Feedback control of LMC contractility

Lymphatic contractions can be modulated in an inotropic and/or chronotropic manner by factors such as transmural pressure, lymph flow, as well as neural and humoral influences [16]. The extent of neural and humoral influences is largely independent of the local mechanical signals experienced by the lymphangion, so is outside the scope of the model described here. Experiments by Gashev et al. on isolated lymphatic vessels have shown that increasing the axial pressure difference (favorable to flow) causes a reduction in both contraction amplitude (negative inotropic effect) and contraction frequency (negative chronotropic effect) [17]. In these experiments, rat mesenteric vessels were exposed to a range of imposed axial pressure differences (0 to 7cmH_2_O) to vary flow (and wall shear stress) through the isolated lymphatic vessel while maintaining the transmural pressure (5cmH_2_O). Because lymphatic flow is largely viscous (Reynolds numbers < 10) and quasi-steady (Womersley parameters < 1), the flow rate and wall shear stress can be assumed to be proportional to and in phase with the imposed axial pressure difference. Gashev et al. observed no significant changes in diastolic diameter, but the systolic diameter increased with axial pressure difference [17]. Contraction amplitude fell more than twofold as the axial pressure difference increased from zero to 3cmH_2_O, showing that there is flow-dependent inhibition of the active lymph pump. The shear stress sensing mechanism is most likely the lymphatic endothelial cell (LEC) surface mechanosensory receptors present at cell-cell junctions (adherens junctions) which cause VEGFR3 phosphorylation [18]. Flow-induced wall shear stress at the endothelial surface induces the production of the vasodilator nitric Oxide (NO) by eNOS in LECs, inhibiting IP3 receptor channels and limiting Ca^2+^ influx into LMCs [19]. Experiments involving inhibiting eNOS (removing basal NO) showed a significant increase in contraction amplitude without increasing contraction frequency, although eNOS inhibition did not completely abolish the effects of flow [17,20]. Only pharmacological blockade of NO and histamine production completely eliminates flow-dependent relaxation of lymphatic vessels [21]. Higher levels of NO production stimulated by acetylcholine (Ach) caused vessel dilatation, decreased tone and decreased contraction frequency. NO acts through soluble guanylate cyclase (sGC) and protein kinase G (PKG) to inhibit IP_3_ receptor channels, limiting Ca^2+^ influx and can also activate SERCA and BKCa channels to increase Ca^2+^ outflux [22,23]. NO induces the hyperpolarization of the lymphatic muscle cells by the activation of ATP sensitive K^+^ channels via cyclic GMP [24]. cGMP/PKG inhibitors reduce the sensitivity to flow induced shear stress and the reduction of lymphatic contractility in the thoracic duct [25]. Other lymphatic endothelium-derived factors (EDFs) such as histamine and prostanoids [16] also play a role in the regulation of contractility by flow [21,26].

In the arterial system, resistance vessels respond to increased luminal pressure by increasing their contractility. This process is mediated through the presence of stretch-activated calcium channels [27]. A similar mechanism is present in lymphatic vessels. In further experiments by Gashev et al., isolated rat mesenteric vessels were exposed to a range of transmural pressures from 1 to 7cmH_2_O. Increasing the luminal pressure without changing the axial pressure difference increased both the frequency and the force of contractions [28,29]. At higher transmural pressures (7-9cmH_2_O for rat mesenteric vessels), lymphatic vessels showed reductions in contraction amplitude [28,30,31]. There is a sharp transition in passive lymphatic vessel stiffness at pressures above 5cmH_2_O due to the arrangement of collagen and elastin fibers [32]; however, the effects of stretch on LMC calcium transients have not been elucidated. Observations in rat thoracic duct show that calcium amplitude and basal levels of calcium did not significantly change with increasing values of stretch - suggesting that the regulation involves a change in the calcium sensitivity of the contractile elements [33]. There was an increase in pacemaker activity (increase in contraction frequency). However, other observations contradict this claim [34].

Ca^2+^ levels in LMCs are regulated through ion channels in the plasma membrane and the boundary of the sarcoplasmic reticulum. Membrane potentials have been measured in endothelial cells of rat mesenteric vessels using wire-myography, under conditions of simulating varying luminal pressure and with/without channel inhibitors (niflumic acid and thimerosal) [14]. If passive force was increased in a preload experiment, action potentials (APs) occurred more regularly than in unstretched vessels and the contraction frequency increased, as did the force generated during each contraction. AP amplitude did not change significantly. L-type Ca^2+^ channels are important for generating strength of contractions, while T-type Ca^2+^ channels regulate lymphatic muscle membrane potential and contraction frequency [14]. It is generally accepted that the phenomenon underlying pressure-dependent modulation is electrical in nature, driven by APs triggering Ca^2+^ influx through L-type Ca^2+^ channels [35,36]. Anoctamin 1 (Ano1) has been identified as the Ca^2+^ -activated Cl^−^ channel providing excitatory Cl^−^ currents critical for mediating the pressure-sensitive modulation of contraction frequency and is a major component of AP generation in murine lymphatics [34].

Intrinsic pumping mechanisms are intricately regulated by mechanical forces acting on the lymphatic vessel [19,37]. For both wall shear-stress sensing by LECs producing NO and stretch sensing by LECs and LMCs subjected to transmural pressure, Ca^2+^ levels in the LMCs can change rapidly due to calcium influx. The autoregulation of contractions is thought to increase lymph flow in response to tissue oedema or gravitational effects [17,28,30,38].

### 1.2 LMC contraction modulation

Dynamic pressure ramp experiments by Davis, Scallan and co-workers [30,38], with independent control of P_in_ and P_out_ while recording vessel dimensions and mid-lymphangion pressure (P_mid_), provide insight into the ability for lymphangions to adapt to pressure conditions. Outlet pressure elevation tests the response of the isolated lymphangion to conditions that would result from a gravitational load outflow obstruction, or tissue compression [30]. Ramp-wise P_out_ elevation led to progressive vessel constriction, a rise in end-systolic diameter and an increase in contraction frequency. Distention of the output segment (the downstream lymphangion), along with slight constriction of the central lymphangion was also observed. Ramp-wise P_out_ elevation also led to a progressive increase in peak systolic P_mid_, so a greater stroke work was needed to compress the lumen against higher pressures [29,39]. The systolic mid-lymphangion pressure necessarily exceeds the outlet pressure P_out_ for every contractile cycle in the pressure ramp that produces ejection. For half of the 16 rat mesenteric vessels tested, the lymphatic pump failed to eject lymph when the outlet pressure exceeded about 10cmH_2_O. Similar experiments involving inlet pressure elevation [38] increased the luminal pressure, the phasic contraction frequency, and the end-diastolic diameter. Pressure-volume curves also showed an increase in lymphangion contractility.

Previous work on lumped-parameter modeling of lymphatics has incorporated mechanical feedback control. Contarino et al. constructed an Electro-Fluid-Mechanical Contraction model using the FitzHugh-Nagumo oscillator to model the lymphangion APs with fluid-mechanical feedback of wall shear stress and wall stretch, and a nonlinear pressure/area relation to simulate the relaxed and constricted lymphatic vessel wall [40]. However, the model only considered the modulation of contraction frequency, not contraction amplitude. Kunert et al. constructed a two-oscillator model coupling the Ca^2+^-mediated contractions triggered by vessel stretch with Nitric Oxide (NO)-mediated relaxation produced by fluid shear stress [41]. The group used the lattice-Boltzmann method to model the spatial distribution, and further mathematical analysis was subsequently published. However, the observations on which the model was based suggest that the inhibition of eNOS leads to loss of lymphatic tone, decreased contraction strength, and increase in contraction frequency [42]. This is contrary to other experiments on eNOS inhibition [43–45] which show an increase in lymphatic tone. Bertram et al. [45] explored the inhibition of contraction strength and frequency by WSS in a single-lymphangion model using three constants (shear stress, delay time, and contraction amplitude) to fit experimental data [17]. The model was then qualitatively compared with existing experimental data performed on rat thoracic duct lymphatic vessels [54]. The effect of wall shear stress was modelled in a ‘black-box’ fashion, where the intracellular machinery of the muscle and endothelial cells lining the lymphangion is not directly represented.

Many experiments and studies have explored how tissue properties and fluid mechanics affect lymph transport [17,28], however less is known about the mechanobiological control mechanisms (active coupling of mechanical signals to biochemical pathways) in lymphatic vessels. There is specifically a lack of quantitative data on the molecular signaling necessary to construct a representative mechanobiological model of the lymphatics.

A multiscale computational model recently developed by our group [46] captures the phasic and tonic contractions in lymphatic muscle and incorporates the subcellular mechanisms within the muscle (phasic and tonic contractile elements, actin-myosin cross-bridge formation, intracellular calcium concentration etc.). This model uses the sliding filament mechanism [47] as adapted for smooth muscle [48], and is integrated within previously validated lumped-parameter models of lymphatic pumping [49–52]. The particular arrangement of the phasic and tonic contractile elements (CEs), with the phasic spring in parallel to phasic CEs and the tonic dashpot in parallel to tonic CEs (Fig. 1D), was the only configuration that produced physiological results out of the many trialed. The phasic spring represents physiologically the elasticity of titin while the tonic dashpot represents the effects of a combination of tonic contractile machinery and the fluid environment around the smooth muscle components. The model provides, for the first time, estimates of the energetics and the efficiency of individual LMCs, and their dependence on pressure conditions.

**Fig. 1.**
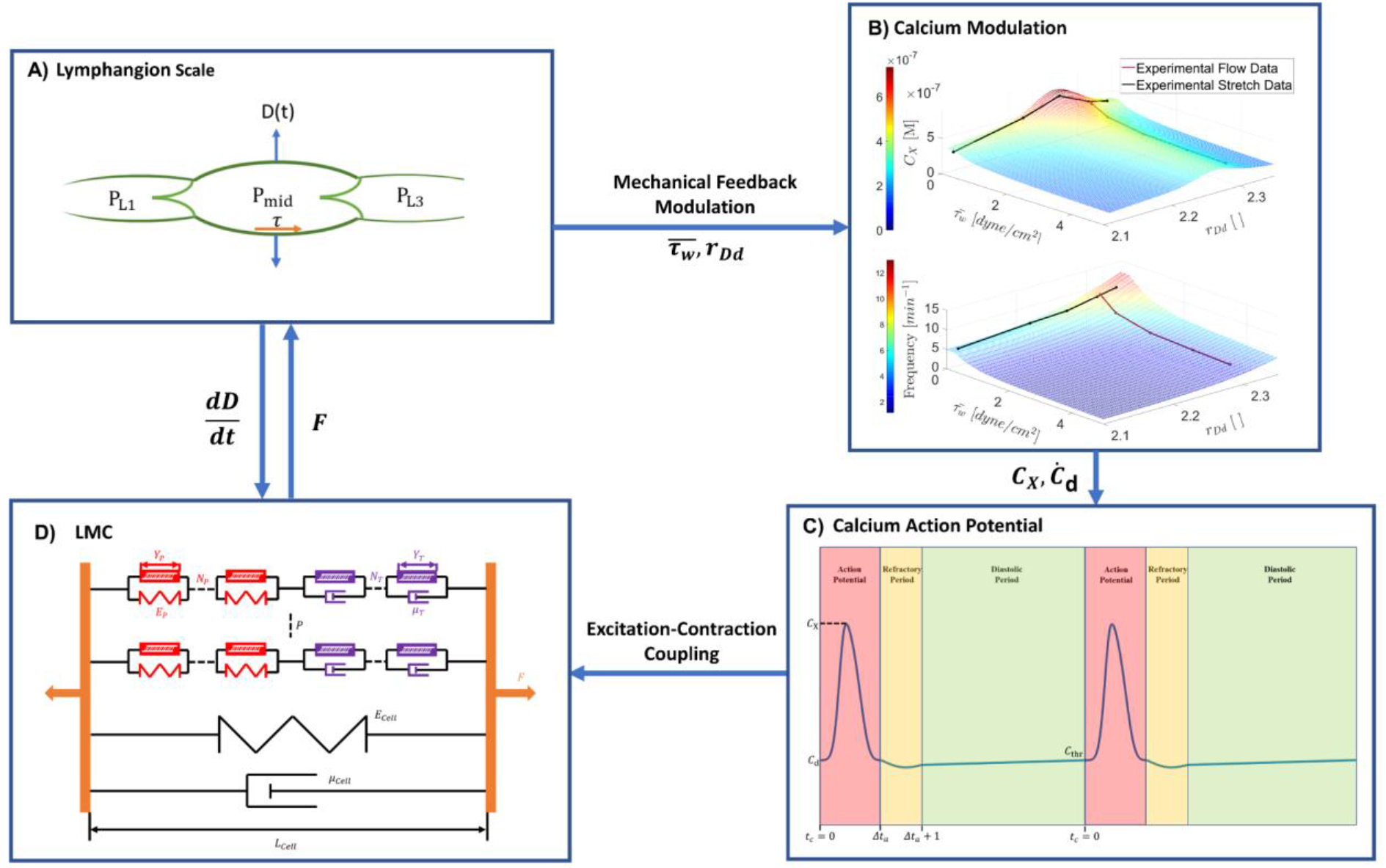
^1^Schematic for the coupling of scales in the muscle model. **A)** Lymphangion scale: the meso-scale model is based on a lumped-parameter model of lymphangion pumping, yielding pressure-flow relations. **B)** Systolic peak calcium and contraction frequency modulation **C)** Calcium waveform of intracellular calcium concentration across two contraction potentials. The periodic waveform consists of three phases: the systolic phase (rapid and short-lived increase and decay of calcium concentration), the refractory phase (period of decrease in calcium concentration, during which subsequent systoles cannot be generated) and the diastolic phase (steadily increasing calcium concentration) **D)** Cellular scale: lymphatic muscle cell contractile force is calculated by the model consisting of two types of contractile elements, with a spring in parallel with phasic elements and a dashpot in parallel with tonic elements. Parallel viscoelasticity represents cell properties. Molecular force generation by contractile elements is calculated from the sliding-filament model (Based on Morris et al., [46]).

The model developed by Morris et al. [46] can be improved by incorporating mechanobiological control feedback to represent more closely the physiological behavior of the lymphangion. The goal of this study was to construct and validate a mechanobiological control feedback model of lymphatics based on experimental data and observations, to understand better the effect of mechanical signals on lymph pumping at a sub-cellular level. The modeling work presented here will further explore the effect of mechanical modulation on the energetics, as understanding this could be useful in characterizing potential treatments.

## 2 Methods

### 2.1 Glossary

|  |  |
| --- | --- |
| **<u>Lymphangion</u>** |  |
| $\mu$ [dynes/cm<sup>2</sup>] | Lymph viscosity (0.01 [dyn/cm<sup>2</sup>]) |
| $L_v$ [cm] | Lymphangion length (0.3 [cm]) |
| $P_{mid}$ [cmH<sub>2</sub>O] | Central lymphangion pressure |
| $P_a$ [cmH<sub>2</sub>O] | Inlet pressure |
| $P_b$ [cmH<sub>2</sub>O] | Outlet pressure |
| $\Delta P_{ab}$ [cmH<sub>2</sub>O] | Axial pressure difference |
| $\Delta P_{tm}$ [cmH<sub>2</sub>O] | Transmural pressure |
| $P_{L1}$ [cmH<sub>2</sub>O] | Passive upstream lymphangion pressure |
| $P_{L3}$ [cmH<sub>2</sub>O] | Passive downstream lymphangion pressure |
| $D_{L1}$ [cm] | Diameter of upstream lymphangion |
| $D_{L3}$ [cm] | Diameter of downstream lymphangion |
| $D(t)$ [cm] | Lymphangion diameter |
| $R_{up}$ [g/cm<sup>4</sup>s] | Upstream cannula resistance |
| $R_{dn}$ [g/cm<sup>4</sup>s] | Downstream cannula resistance |
| **<u>Calcium</u>** |  |
| $c(t)$ [M] | Intracellular calcium concentration |
| $C_x$ [M] | Peak intracellular calcium concentration |
| $\dot{C}_d$ [s<sup>-1</sup>] | Gradient of calcium concentration in the diastolic phase |
| $\dot{C}_r$ [s<sup>-1</sup>] | Gradient of calcium concentration in the refractory phase |
| $t_c$ [s] | Time into a contractile cycle |
| $\Delta t_a$ [s] | Systole time |
| $C_0$ [M] | Peak intracellular calcium concentration below threshold shear stress ($\tau_{thr}$) |
| $m_{C\tau}$ [M] | Modulation of $C_x$ due to $\tau_w(t)$ |
| $m_{CD}$ [ ] | Modulation of $C_x$ due to $r_{Dd}$ |
| $NO$ [ ] | Normalized production of NO [0-1] induced by $\tau_w(t)$ |
| $m_{fD}$ [ ] | Stretch Factor - modulation of contraction frequency induced by diastolic stretch ratio $r_{Dd}$ |
| **<u>Energetics</u>** |  |
| $\eta_1$ [%] | Efficiency of LMCs in doing work |
| $\eta_2$ [%] | Efficiency in producing lymph flow |
| $E_P$ [erg] | Energy liberated by phasic contractile elements |
| $E_T$ [erg] | Energy liberated by tonic contractile elements |
| **Mechanical** |  |
| $\tau_w(t)$ [dyne/cm<sup>2</sup>] | Wall shear stress on lymphangion |
| $\tau_{thr}$ [dyne/cm<sup>2</sup>] | Threshold shear stress for modulation |
| $r_{Dd}$ [ ] | Diastolic stretch ratio |
| $r_{Dd1}$ [ ] | Upstream diastolic stretch ratio |
| $r_{Dd3}$ [ ] | Downstream diastolic stretch ratio |

### 2.2 Lymphangion model

The lumped-parameter model we used were based on models developed by Bertram et al. [2,45,49–51]. Extensive details of the equations and method of solution used are available in those papers and their appendices.

To facilitate the replication of experiments performed on rat mesenteric lymphatic vessels, passive (not actively contracting) upstream and downstream lymphangions have been included (represented by pressures P_L1_ and P_L3_ and resistances R_L1_ and R_L3_) (Fig. 1a). Although the upstream and downstream lymphangions do not contract, we have incorporated their pressure signals into the control algorithm to test for the potential importance of signaling between adjacent lymphangions (see sections 3.3 and 3.4). Electrical coupling between adjacent lymphangions occurs via gap junctions [15,53] and between muscle cells in the lymphatic wall [44]. Similar conduction speeds were found for electrical coupling between muscle cells within a lymphangion and electrical coupling across valves driving inter-lymphangion coordination, suggesting similar mechanisms [15]. The timing difference between contractions in adjacent lymphangions is usually less than 0.5s, and commonly less than 0.25s but there are large variations.

Upstream and downstream valve resistances (R_V1_ and R_V2_) have a transition in resistance from low forward-flow resistance to high backflow resistance. The valve resistance is modelled as a double sigmoidal function, where small positive pressure differences across the valve causes a decrease in the resistance due to the opening of the valve [49].

Cannula resistances (represented as resistances R_up_ and R_dn_) are included to replicate the conditions of experiments performed on isolated rat mesenteric lymphatics. The cannula resistances are calculated using Poiseuille’s equation. Mass conservation in the model was verified by comparing the cycle-average inlet and outlet flow rates. These values were within 1% of each other for all results shown.

### 2.3 Muscle cell model

We have adapted an existing multi-scale model of lymphatic muscle cells (LMCs) [46] based on a combination of the molecular sliding filament model of Huxley and its adaptations for smooth muscle. The model consists of three scales: molecular, cellular, and lymphangion. There are two types of contractile elements connected in series, with a strain-stiffening spring in parallel to the phasic elements and a Newtonian dashpot in parallel with tonic elements. Effects of excitation-contraction coupling (ECC) are included at the molecular scale to induce periodic contractions (Figs. 1c and 1d). Detailed description of the equations in the model describing phasic and tonic forces, crossbridge head displacements, attachment rate and saturation functions, and energetics is given by Morris et al. [46]. The partial differential equations for the displacement-distribution of myosin heads (for both phasic and tonic CEs) were solved with a second-order Godunov scheme [46].

### 2.4 Molecular scale modeling

The molecular models of force generation by phasic and tonic contractile elements (CEs) are based on Huxley’s sliding-filament model [47] and its adaptation for smooth muscle [48]. The equations of these molecular models are detailed by Morris et al. [46]. In this paper, we study the effects of mechanical signals on lymphatic pumping, and more specifically on LMC behaviour. Therefore, we are interested in how the phasic and tonic forces, LMC efficiency (eq. (1)), and pump power (eq. (2)) change with mechanical signaling.

The efficiency of LMCs in doing work (*η*_1_) is calculated as the ratio of the total work done by LMCs to the total energy liberated (the sum of energy liberated by phasic and tonic CEs). Liberated energy not used to do work is lost as heat energy [46].

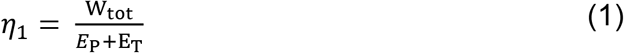

Efficiency in producing lymph flow (*η*_2_) is calculated as the ratio of the pump power produced by the lymphangion to the total energy liberated. Note that in simulations of positive axial pressure difference, efficiency to lymph transfer becomes negative, as passive flow becomes a larger driving force for lymph flow than active pumping.

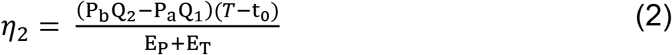

where (*T* − t_0_) is the total duration of a contractile cycle, 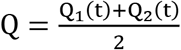, with Q_1_ and Q_2_ being the mean flow rates through the inlet and outlet valves respectively over the contractile cycle, P_b_ is the outlet pressure, and P_a_ is the inlet pressure.

#### 2.4.1 Excitation-contraction coupling

Intracellular free calcium ion concentration *c*(*t*) regulates the contractions of both striated and smooth muscle types. Here, *c*(*t*) is then used to calculate the saturation of binding proteins, which alters the transition rates between myosin head states, influencing force generation [46]. The input calcium transient waveform was based on rat mesenteric lymphatic vessel measurements of Zawieja et al. [12], which showed a systolic peak of 232 nM and a diastolic plateau of 140 nM (Fig. 1c). The mathematical expression is:

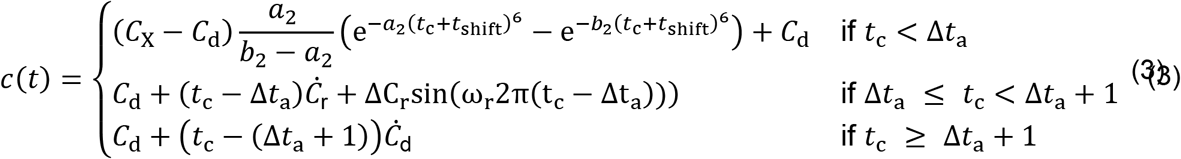

If *C*_X_ ≥ *C*_thr_ and *t*_c_ ≥ Δ*t*_a_ + 1, reset *t*_c_ = 0 and start the next calcium systole.

In the above equation (Eq. (3)), C_X_ is the peak calcium concentration, C_D_ is the basal diastolic calcium concentration, *a*_2_ and *b*_2_ are constitutive parameters to fit the calcium trace [12], t_c_ is the time from the beginning of the current cycle, Δt_a_ is the time in the current cycle from the beginning of the refractory phase, and ΔC_r_ is the extent of the decrease in calcium concentration during the refractory phase. The refractory phase is modelled as a negative sinusoidal function superimposed onto a line with a negative gradient Ċ _r_. No subsequent calcium systole can be initiated during the refractory phase of a cycle. Following the refractory phase is the diastolic phase, a line with positive slope Ċ_d_. A new calcium systole is initiated once the calcium concentration reaches the value C_thr_. The intracellular calcium concentration at a specific time point is used to compute the saturations of the regulatory proteins TnC and CaM [46].

In order to introduce the effects of mechanical signals on the lymphangion, the parameters C_X_ (the systolic peak value of calcium concentration), and Ċ_d_ (the gradient of the diastolic phase) are modulated by mechanical signals. Changing C_X_ creates an inotropic effect (modulating contractile force), whereas changing Ċ_d_ creates a chronotropic effect (modulating contraction frequency).

The model does not directly model the voltage-dependent calcium channels (VDCCs) and their respective ion fluxes, as much of the identity and importance of different ion channels in response to mechanical signals is not well characterized. With the focus on regulation of mechanical events by stretch and fluid shear, it enables us to avoid the complexities of electrochemical oscillator modelling, while allowing us to link muscle action to calcium concentration changes.

### 2.5 Mechanical signals

#### 2.5.1 Flow-induced wall shear stress

The spatially averaged instantaneous wall shear stress τ(t) for the lymphangion is computed as:

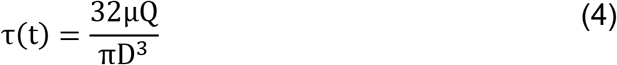

where μ is the lymph viscosity, Q is the mean flow rate. D is the lymphangion diameter. The cycle time average wall shear stress (TAWSS) of the lymphangion 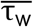 is used to modulate the contractility of the lymphangion in the subsequent cycle:

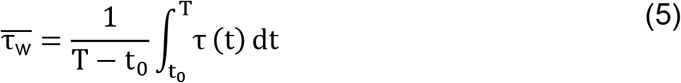

where t_0_ is the start and T is the end time of the contractile cycle.

#### 2.5.2 Pressure-induced stretch

The feedback signal for pressure-induced stretch is based on the time-average diastolic of the previous cycle. We define stretch (r_Dd_) as the time average diastolic diameter 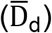 divided by the diameter at zero transmural pressure, D_d0_. Diastole is defined as the point at which:

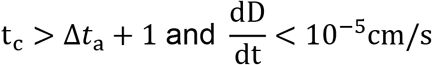

Two alternative stretch control signals were tested, based on stretch signaling from the neighboring passive upstream and downstream lymphangions (with associated pressures P_l1_ and P_l3_). This corresponds to the propagation of pressure-derived information amongst adjacent lymphangions. The alternative stretch ratios are:

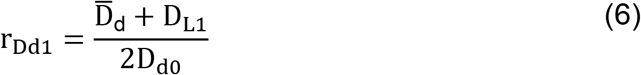

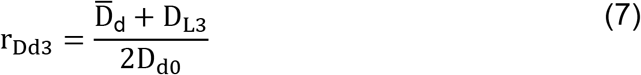

where D_L1_ and D_L3_ are the cycle-average diameters of the passive upstream and downstream lymphangions respectively.

#### 2.5.3 Averaging algorithm

Cycle-average wall shear stress 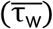 and pressure-induced stretch r_Dd_ over the contractile cycle that has just been completed modulate calcium parameters C_X_ and Ċ_d_ for the next contractile cycle. Therefore, the intracellular calcium amplitude and contraction frequency of the next contractile cycle are determined upon completion of the current contractile cycle. We also considered an instantaneous algorithm, in which the mechanical signals at a specific time point instantaneously modulate the current value of: C_X_ and Ċ_d_. However, because the calcium transient precedes both contraction and any subsequent change in lymph flow (due to the time delay/phase difference between the calcium oscillation and the formation of actin-myosin cross-bridges), instantaneous control was determined to be not physiologic.

### 2.6 Mechanical feedback modulation

Since we will need to refer to pressure differences frequently, we define the axial pressure difference *P*_a_ − *P*_b_ as Δ*P*_ab_, noting the this value is positive for a pressure gradient favourable to forward flow, and the transmural pressure *P*_mid_ − *P*_ext_ as Δ*P*_tm_, noting that this is positive for a value producing vessel distension. Parameters of the modulation are matched to experimental data performed on rat mesenteric lymphatic vessels [17,28]. The parameters for shear stress modulation are based on experiments of Gashev et al. (2002), where the axial pressure difference was varied while keeping the transmural pressure constant at 5cmH_2_O for all simulations. The parameters for stretch modulation are based upon the experiments of Gashev et al. (2004), where the transmural pressure was varied while the axial pressure difference is held at 0 cmH_2_O for all experiments.

#### 2.6.1 Intracellular peak calcium modulation

The mechanical signals of flow-induced shear stress and pressure-induced stretch modulate the peak intracellular calcium concentration C_X_:

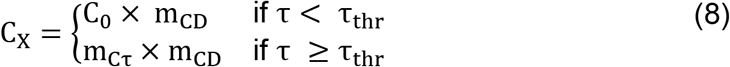

where C_0_ is the calcium concentration at zero axial pressure difference, m_Cτ_ is a function of wall shear stress and m_CD_ is a function of stretch.

Using existing equations and parameters in this sub-cellular lymphangion model, peak intracellular calcium concentrations were matched with the corresponding normalized systolic diameters quoted in [17] (Fig. 2c). The five data points in the plot represent the cycle-average wall shear stress experienced by the model lymphangion by applying positive axial pressure differences of 0, 1, 3, 5, and 7cmH_2_O (the same pressure differences applied to the experimental lymphangion)^2^. A power function (eq. (9)) provided a good fit to the experimental data (*r*^2^= 0.998).

**Fig. 2.**
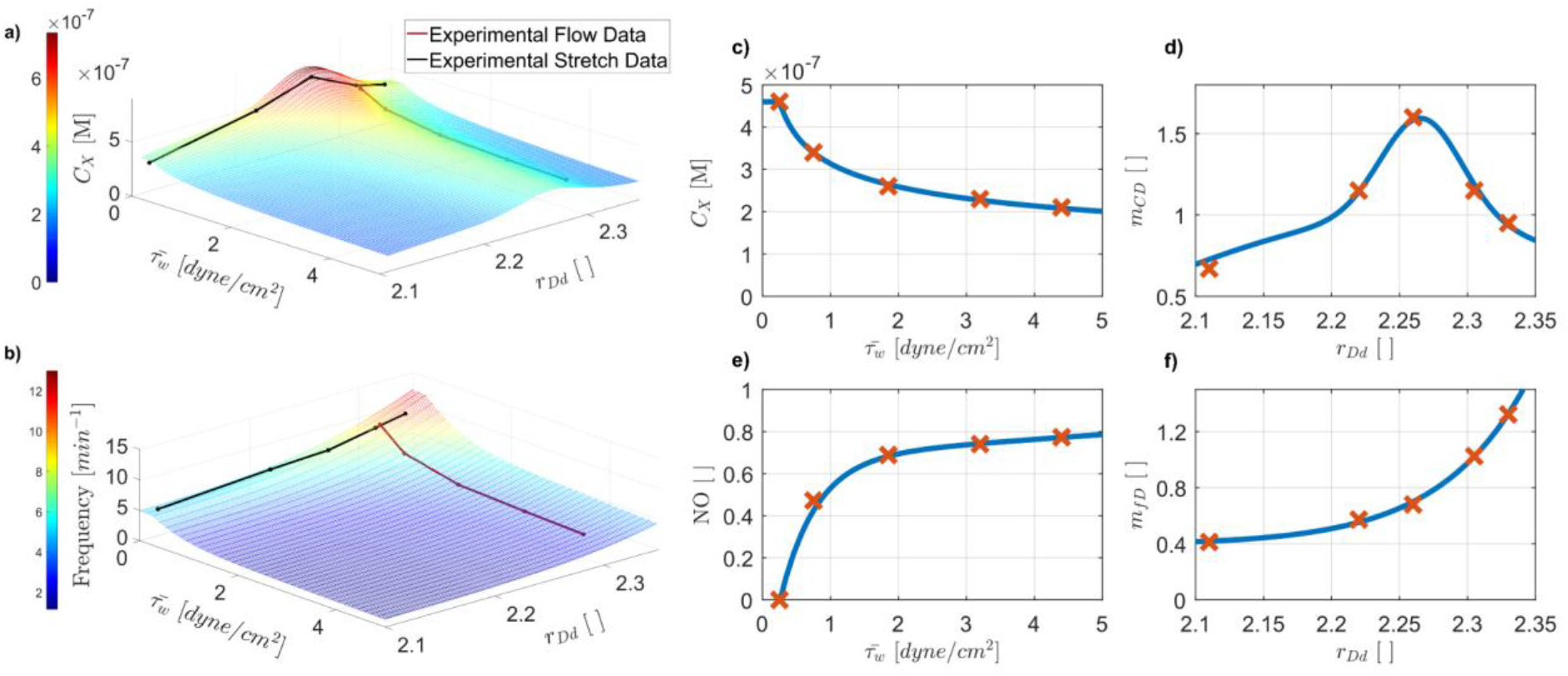
Mechanical feedback modulation functions. **(a)** 3D plot of peak intracellular calcium modulation (combining curves **(c)** and **(d)**). **(b)** 3D plot of contraction frequency (combining curves **(e)** and **(f)**). Contraction frequency is related to the gradient of the diastolic period (*Ċ*_d_). Blue curves **(c, d, e, f)** show the modulation of parameters calculating **(c, d)** intracellular calcium peak *C*_*X*_ (*m*_*Cτ*_, *m*_*CD*_) and **(e, f)** gradient of the diastolic period (*Ċ*_d_) (*m*_*fD*_ and NO). Red data points **(c, d, e, f)** indicate the predicted intracellular calcium peak and contraction frequency to match the macroscopic behaviour of lymphangions [17,28].

**Fig. 3.**
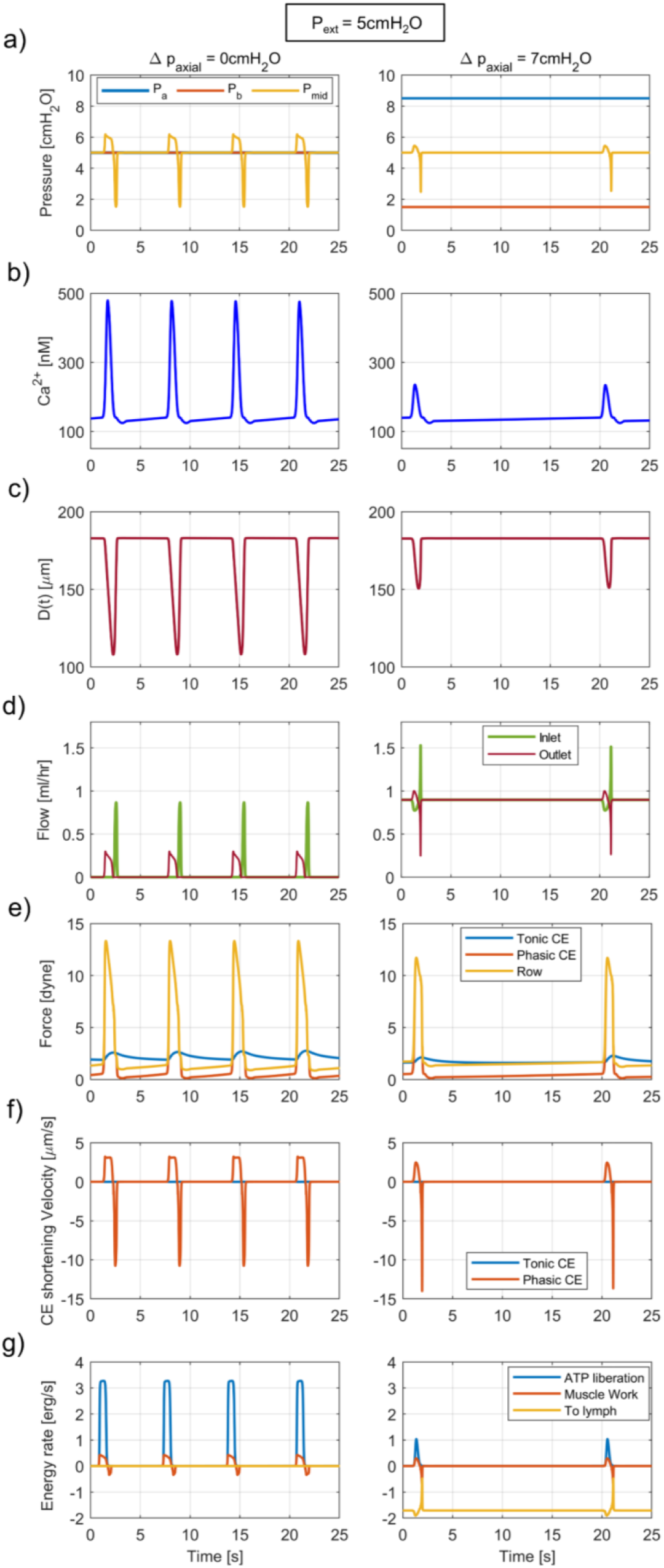
Plots showing the effect of shear on lymphangion parameters and calcium under conditions of Δ*P*_ab_ = 0cmH_2_O (left panels) and 7cmH_2_O (right panels), with transmural pressure fixed at 5cmH_2_O. **(a)** Inlet pressure P_a_ increased, and outlet pressure P_b_ decreased by the same amount to produce constant Δ*P*_ab_ of 0 and 7cmH_2_O. Mid-lymphangion pressure P_mid_ increases during systole to expel fluid, followed by a decrease to refill the lymphangion via the suction effect. **(b)** Intracellular calcium concentration. **(c)** Central lymphangion diameter. Diastolic diameter displays no change whereas systolic diameter increases due to increased flow. **(d)** Inlet and outlet flow rates. Increased Δ*P*_ab_ increases mean flow rate. In the 0cmH_2_O case, systolic contractions increase outlet flow rate, followed by an increase in inlet flow rate due to the suction effect. In the 7cmH2O case, inlet and outlet flow rates counteract as both inlet and outlet valves are open with positive axial pressure. Both valves are open for the entire simulation at this Δ*P*_ab_. **(e)** Force contributions from the subcellular contractile elements: phasic, tonic, and total row forces. The phasic CE (orange) contributes significantly to the cell force during systole, whereas the tonic CE (blue) contributes significantly during the diastolic phase. There is a decrease in both phasic and tonic forces due to increased flow. **(f)** Shortening velocity of the CEs. The phasic CE shortening velocity increases during systolic contraction, followed by a lengthening velocity (negative shortening velocity) to distend the lymphangion due to the suction effect. The small tonic CE shortening velocity is due to the tonic dashpot, resisting significant changes in length in the tonic CE **(g)** Rate of useful work done (orange) in comparison to the energy liberated by ATP hydrolysis (blue). Energy transferred to lymph flow (yellow) is also calculated. This value is negative for favorable Δ*P*_ab_ as the pressure difference becomes the main driving force.

To match the normalized systolic diameters quoted in [28] (Fig. 2d), a multiplying factor m_CD_ modulates the intracellular systolic calcium concentration. The five data points in the plot represent the results from imposing transmural pressures of 1, 2, 3, 5, and 7cmH_2_O respectively [28]. There is an increase in diastolic diameter with increasing values of transmural pressure, due to the passive pressure-diameter relationship of rat mesenteric lymphatic vessels [32]. Maximum contraction amplitude was observed at a transmural pressure of 3cmH_2_O, with a diameter change of greater than 41.5% (diameters are normalized by the diameter in Ca^2+^ free APSS). A double Gaussian curve fit (Eq. (10)) provided the best fit to the experimental data (*r*^2^ = 0.996).

##### 2.6.1.1 Effect of wall shear stress on peak calcium

Modulation curve for m_Cτ_:

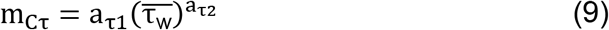

m_Cτ_ is a power function (Fig. 2c), where increasing values of cycle-average wall shear stress in the preceding contractile cycle lead to a decrease in intracellular peak calcium C_X_ in the subsequent contractile cycle.

A threshold for shear stress modulation of peak calcium is included to prevent possible numerical oscillations in peak calcium between contractile cycles. It has been shown in rat thoracic duct [54] that there is a minimum wall shear stress τ_thr_ for inhibition of the contractility of rat mesenteric vessels. Below τ_thr_, the peak intracellular calcium is constant at 4.60 ×10^−7^ [M] (Fig. 2c). In the model this value is the cycle-average wall shear stress 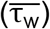 when there is no imposed axial pressure difference, and 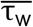 purely arises from shear stress due to contractions [45].

##### 2.6.1.2 Effect of stretch on peak calcium

Modulation curve for m_CD_:

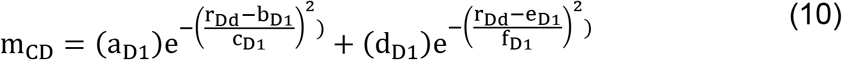

The modulation curve for the stretch multiplication factor m_CD_ as a function of the stretch ratio r_Dd_ is a double Gaussian. This function m_CD_ is multiplied by either *C*_0_ if 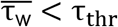 or m_Cτ_ if 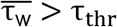, to modulate the intracellular peak calcium C_X_ (Fig. 2d). For rat mesenteric vessels, increasing values of stretch causes an increase in contraction peak, up to a certain level of stretch (3cmH_2_O transmural pressure), then the contraction amplitude falls with stretch corresponding to transmural pressures of 5 and 7cmH_2_O [28].

#### 2.6.2 Contraction frequency modulation

The contraction frequency of the lymphangion is modulated by varying Ċ_d_, the gradient of the line during the diastolic period. Ċ_d_ is determined by two variables: normalized nitric oxide (NO) production, with increasing NO resulting from increased flow-induced wall shear stress, and m_fD_, resulting from pressure-induced stretch. The value of NO is normalized due to a lack of quantitative measurements of NO concentration in response to the application of shear stress on cells. A normalized approach allows us to model the effect of NO without knowledge of the actual concentrations.

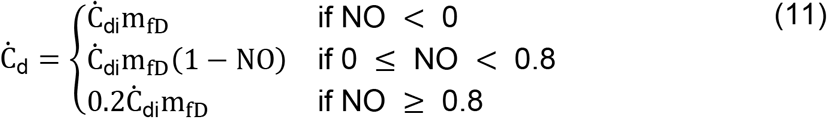

For values of NO below 0, there is no inhibition of contraction frequency. For NO values above 0.8, Ċ_d_ is fixed at the minimum value of Ċ_di_m_fD_0.2 (corresponding to a contraction frequency of 3/min).

In their experiments with varied axial pressure difference, Gashev et al. observed significant flow-dependent inhibition of the frequency and amplitude of lymphatic contractions (negative chronotropic and inotropic effects). The results showed that contraction frequency decreased as favorable axial pressure difference increased (eq. 12 and Fig. 2e). A double exponential function provided the best fit to the experimental data (*r*^2^ = 1.000). Mesenteric lymphatics significantly increased contraction frequency upon raising transmural pressures, providing a 1.8-fold increase between 1 and 7cmH_2_O [28]. Modulation of Ċ d by stretch is shown in eq. 13 and Fig. 2f. A power law function provided the best fit to the experimental data (R-square = 0.998).

##### 2.6.2.1 Effect of wall shear stress on contraction frequency

Modulation curve for NO production:

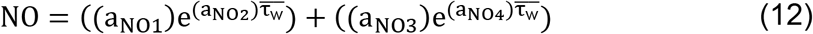

NO is a double exponential function (Fig. 2e), where increasing values of cycle-average wall shear stress lead to a larger value of NO, reducing the value of Ċ_d_, the gradient of the diastolic phase of the cycle, and therefore increasing the time period for a contractile cycle. Inhibition of contraction frequency can only occur at values of wall shear stress exceeding τ_thr_.

##### 2.6.2.2 Effect of stretch on contraction frequency

Modulation curve for the stretch factor (m_fD_) which modulates the contraction frequency:

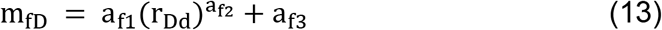

The modulation curve for m_fD_ is a power-law function (Fig. 2e), where increasing values of stretch (r_Dd_) lead to a larger value of m_fD_.

### 2.7 Mechanobiological feedback as function of shear and stretch

The modulation curves from sections 2.6.1 (peak calcium concentration modulation) and 2.6.2 (contraction frequency modulation) are combined in Figs. 2a and 2b to form surfaces to visualize the overall effect of mechanical signals on the active pumping of lymphangions. The x and y axes are independent variables for the mechanical signals of flow-induced wall shear stress and pressure-induced stretch. The z-axis in Fig. 2a represents the peak systolic calcium concentration as a function of the two mechanical signals, and the z-axis in Fig. 2b similarly represents the contraction frequency.

Red lines on the surfaces indicate results the imposed axial pressure difference experiments [17], while black lines indicate results from the transmural pressure experiments [28]. There are limited experimental data on rat mesenteric lymphatic vessels, and the surfaces may be unreliable in regions far from available experimental data. However, these surface fits are a first step to gain an overall understanding of how active pumping of lymphangions is modulated by mechanical signals at a subcellular scale when considering their different muscle types (phasic and tonic) and behaviors at a sub-cellular level (cross-bridge formation, calcium concentration).

#### 2.7.1 Systolic peak calcium modulation

The peal calcium surface (Fig. 2a) combines the modulation functions presented in 2.6.1 (Figs. 2c and 2d), to show how the systolic peak calcium is modulated by the two mechanical signals. The surface shows an asymptotic decrease in peak calcium with increasing values of shear, and a Gaussian behaviour with respect to stretch (initial increase until a stretch ratio of 2.27 before a decrease for larger stretch ratios).

#### 2.7.2 Contraction frequency modulation

The contraction frequency surface (Fig. 2b) combines the modulation functions presented in 2.6.2 (Figs. 2e, and 2f), to show how the contraction frequency is modulated by the two mechanical signals. The curve shows an asymptotic decrease in contraction frequency with increasing values of shear and an exponential increase in contraction frequency with increasing values of stretch ratio.

### 2.8 Model Simulations

#### 2.8.1 Model verification simulations

The results of quasi-steady experiments on isolated lymphangions were used in the construction of the model. These experiments were performed with variations in either axial pressure difference or transmural pressure, keeping the other constant. They are thus well suited for the specification of the control parameters (eqs. 8–13). Comparison with the model without the modulation algorithm is also shown (Fig. 4), where C_X_ and Ċ_d_ are fixed at the values corresponding to the TAWSS when Δ*P*_ab_ is zero, and Δ*P*_tm_ = 5cmH_2_O for axial pressure difference experiments or Δ*P*_tm_ = 1cmH_2_O for transmural pressure experiments.

**Fig. 4.**
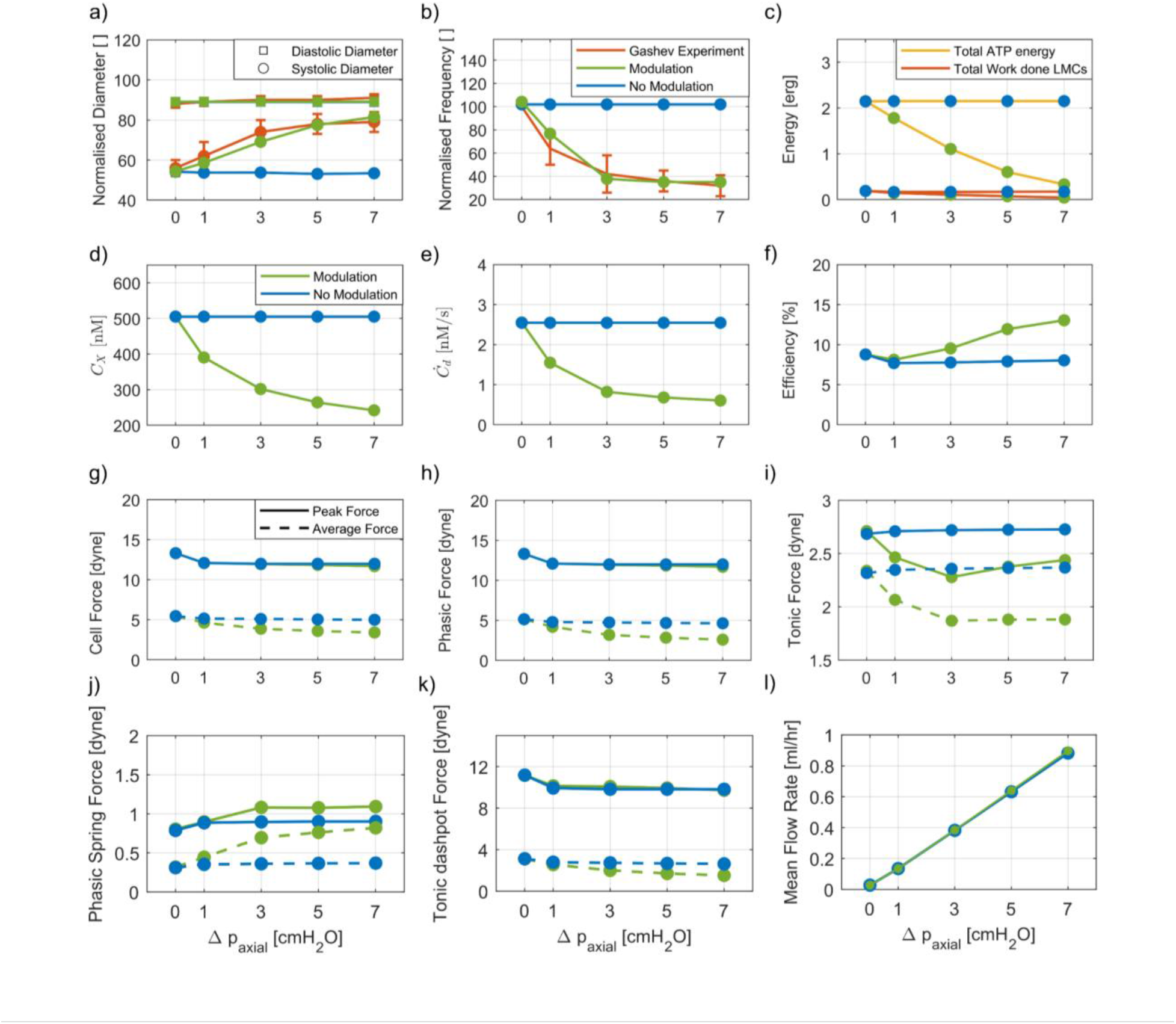
Plots showing the effect of shear on macroscopic lymphangion parameters and calcium. The model lymphangion with modulation (green) is compared to without mechanical modulation (blue). **(a)** Systolic and diastolic diameters as Δ*P*_*ab*_ is varied. Increasing flow increases the lymphatic systolic diameter, while minimally modulating the diastolic diameter in the modulated case (green) and in the experiments of Gashev et al. (red). Unmodulated model response (blue) shows no change in systolic diameter in response to increasing flow. **(b)** Normalized contraction frequency, showing that increasing flow produces strong chronotropic inhibition. Unmodulated model (blue) shows no change in frequency in response to increasing flow. **(c)** Summary of howe total ATP energy and work done by LMCs during a contractile cycle varies with flow. Due to fixed calcium, unmodulated energy values remain constant, whereas both parameters decrease for the modulated model lymphangion. **(d)** Summary of how *C*_*X*_ changes with Δ*P*_*ab*_. The modulated case shows a gradual decrease, whereas unmodulated remains constant. **(e)** How ‘*Ċ*_d_ ‘ (gradient of the diastolic phase) changes with Δ*P*_*ab*_. **(f)** Summary of the effect of flow on the efficiency (*η*_1_) of LMCs in converting energy from ATP into muscle work. The modulated model lymphangion shows an increase in efficiency, whereas efficiency for the unmodulated case remains constant. **(g)** Peak and average cell force (LMC force in a single row). Peak modulated force (green) is less than peak force without modulation (blue). **(h)** Peak and average phasic force. Phasic force increases until LMCs contract and expel lymph through the outlet valve. Peak force is independent of calcium modulation; average phasic force decreases due to decrease in calcium concentration and increase in total time taken to complete the calcium systole. **(i)** Peak and average tonic force. Both decrease with modulation due to decreased calcium. **(j)** Peak and average phasic spring force. Both increases with modulation as result of reduced average phasic CE force **(k)** Peak and average tonic dashpot force. Tonic dashpot force balances force from phasic CEs. **(l)** Cycle-mean flow rate. The modulated mean flow rate is greater compared to the unmodulated mean flow rate, suggesting no advantage of maintaining lymphatic contractions in high flow situations.

**Fig. 5.**
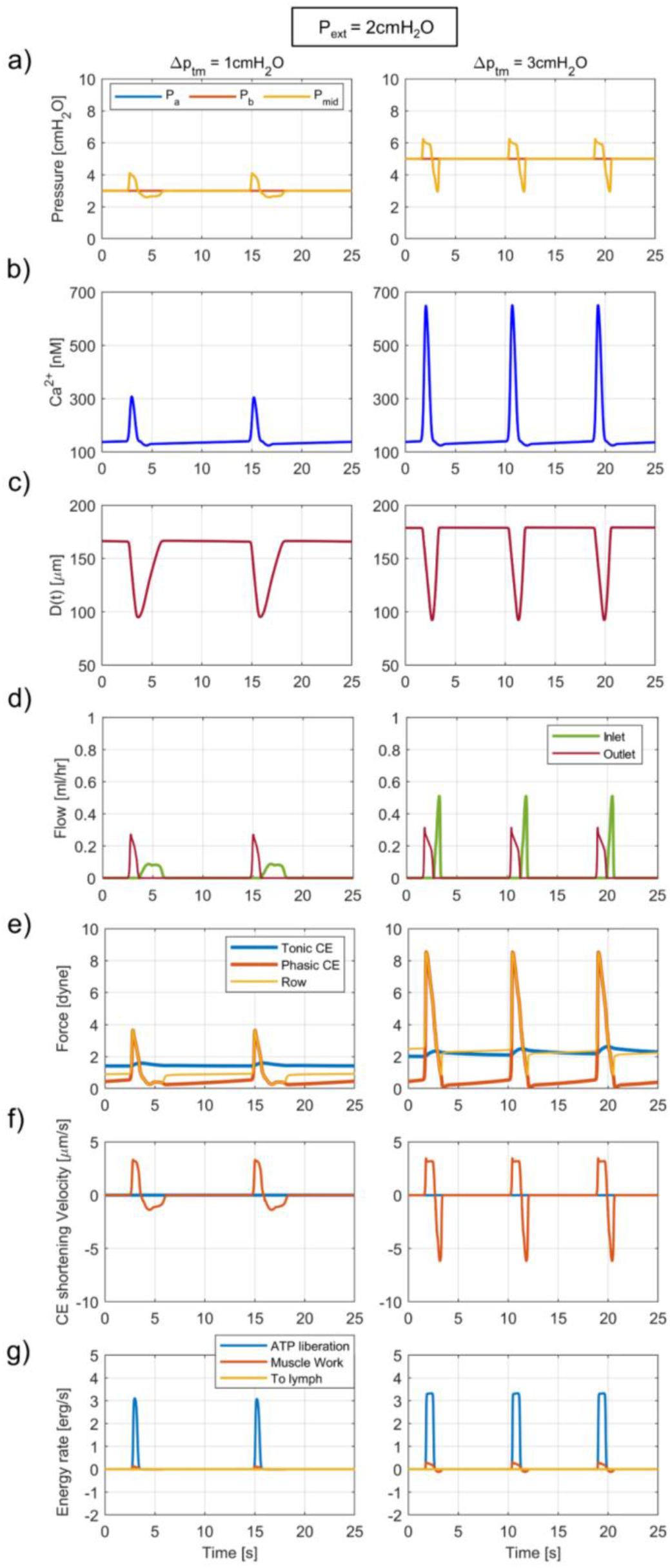
Plots showing the effect of stretch on lymphangion parameters under conditions of 1cmH_2_O (left panels) and 3cmH_2_O (right panes) transmural pressure, with axial pressure fixed at zero. **(a)** Inlet pressure *P*_*a*_ and outlet pressure *P*_*b*_ increased by the same amount to produce to produce transmural pressures Δ*P*_*tm*_ of 1 and 3cmH_2_O. **(b)** Intracellular calcium concentration. Increasing Δ*P*_*tm*_ from 1 to 3cmH_2_O results in increased calcium peak concentration and frequency. **(c)** Diameter of central lymphangion. Diastolic diameter increases with increasing stretch (passive pressure-diameter relationship). Contraction amplitude increases in response to increased calcium. **(d)** Inlet and outlet flow rates. **(e)** Force contributions from the subcellular contractile elements: phasic, tonic, and total row forces. **(f)** Shortening velocity of phasic and tonic CEs. The phasic CE shortening velocity increases during systolic contraction, followed by lengthening (negative shortening velocity) to distend the lymphangion due to the suction effect. The small tonic CE shortening velocity is due to the tonic dashpot. **(g)** Rate of useful work done (orange) in comparison to the energy liberated by ATP hydrolysis (blue), and energy transferred to lymph flow (yellow).

**Fig. 6.**
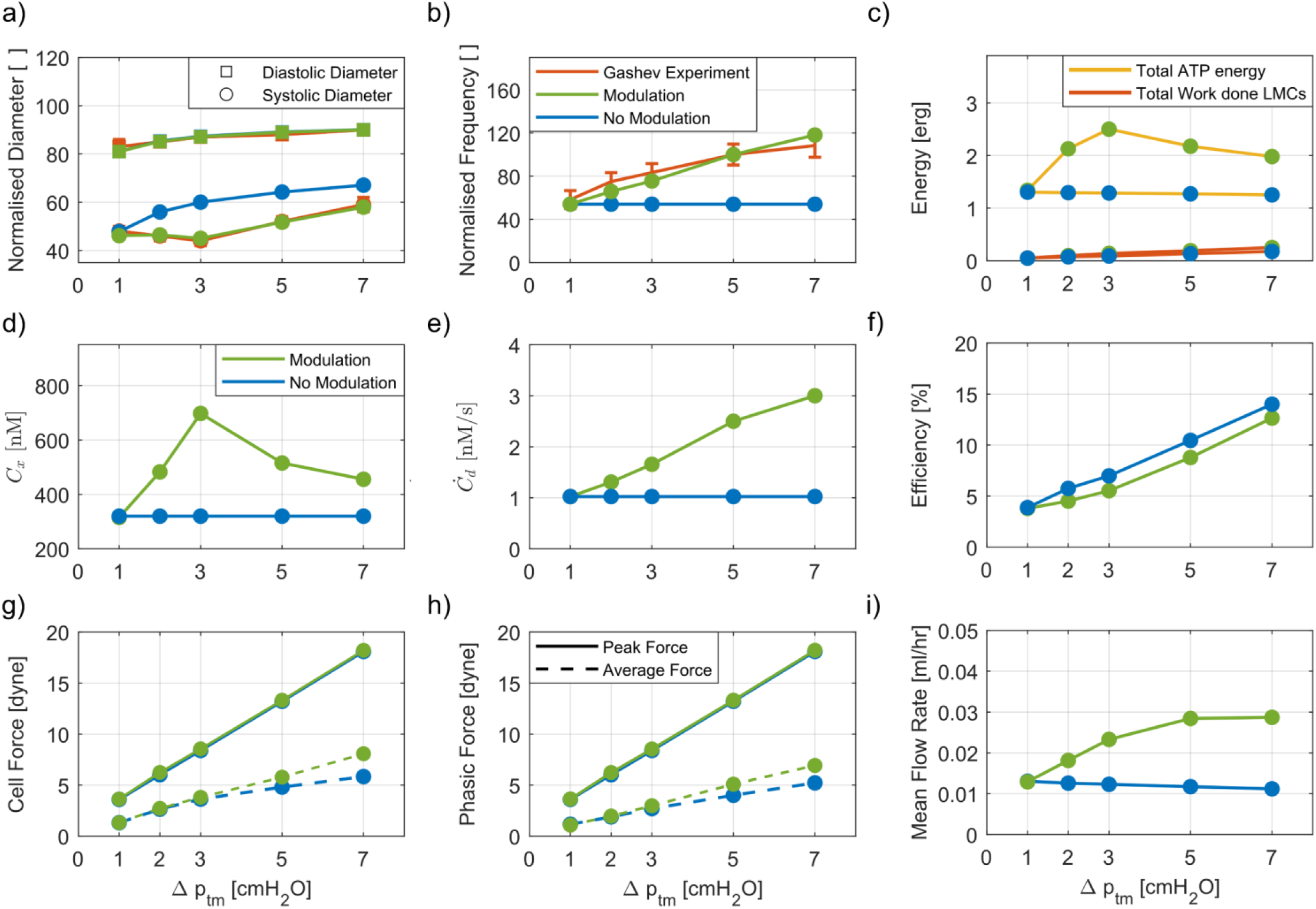
Plots showing the effect of stretch on macroscopic lymphangion parameters and calcium. The model lymphangion with modulation (green) is compared to without mechanical modulation (blue). **(a)** ESD and EDD as Δ*P*_*tm*_ is increased. Increasing Δ*P*_*tm*_ increases the lymphatic diastolic diameter, while decreasing the systolic diameter, followed by an increase in systolic diameter. Unmodulated model response (blue) shows a consistent increase in systolic diameter with increasing stretch. **(b)** Normalized contraction frequency, showing that increasing stretch increases contraction frequency. **(c)** Total ATP energy and work done by LMCs over a single contractile cycle. **(d)** *C*_*X*_ changes with Δ*P*_*tm*_. The modulated case shows gradual increase up to 3cmH_2_O, followed by a decrease, whereas unmodulated remains constant. **(e)** *C*_d_ (gradient of the diastolic phase) changes with Δ*P*_*tm*_. **(f)** Effect of stretch on the efficiency (*η*_1_) of LMCs in converting energy from ATP into muscle work. Both the modulated and the unmodulated model lymphangion show an increase in efficiency in response to increasing stretch. **(g)** Peak and average phasic force **(h)** Peak and average cell force **(i)** Cycle mean flow rate. The modulated mean flow rate is greater compared to the unmodulated mean flow rate. Mean flow rate decreases for the unmodulated case.

#### 2.8.2 Model validation simulations

As a test of the model with both shear stress and stretch modulation incorporated, dynamic pressure ramp experiments [30,38] are used to validate the model. Modulating either the upstream or downstream pressure while keeping the other constant creates simultaneous variations in transmural pressure and axial pressure difference.

##### 2.8.2.1 Afterload ramp simulation

As a test of the model with both shear stress and stretch modulation incorporated, an outlet pressure ramp was assessed in which the outlet pressure P_b_ was ramped from 1cmH_2_O to 12cmH_2_O at a rate of 3.7cmH_2_O/min and the inlet pressure P_a_ was maintained constant at 1cmH_2_O (Fig. 7). P_b_ was then held at 12cmH_2_O for 70s before decreasing back down to 1cmH_2_O. P_ext_ held at constant value of 0cmH_2_O, setting Δ*P*_tm_ = 1cmH_2_O when P_b_ = 1cmH_2_O. P_b_ elevation results in a rise in end-systolic diameter (ESD) and an increase in contraction frequency. Davis et al. [30] concluded that the lymphatic muscle cells intrinsically increase their contractility during an afterload ramp. A higher cannula resistance was used for these simulations: the cannula resistance in the model was raised to 10^8^ g/cm^4^s [38]. Stretch signaling where r_Dd_ modulates the systolic peak calcium and r_Dd3_ modulates contraction frequency produced results that closely match the experimental results.

**Fig. 7.**
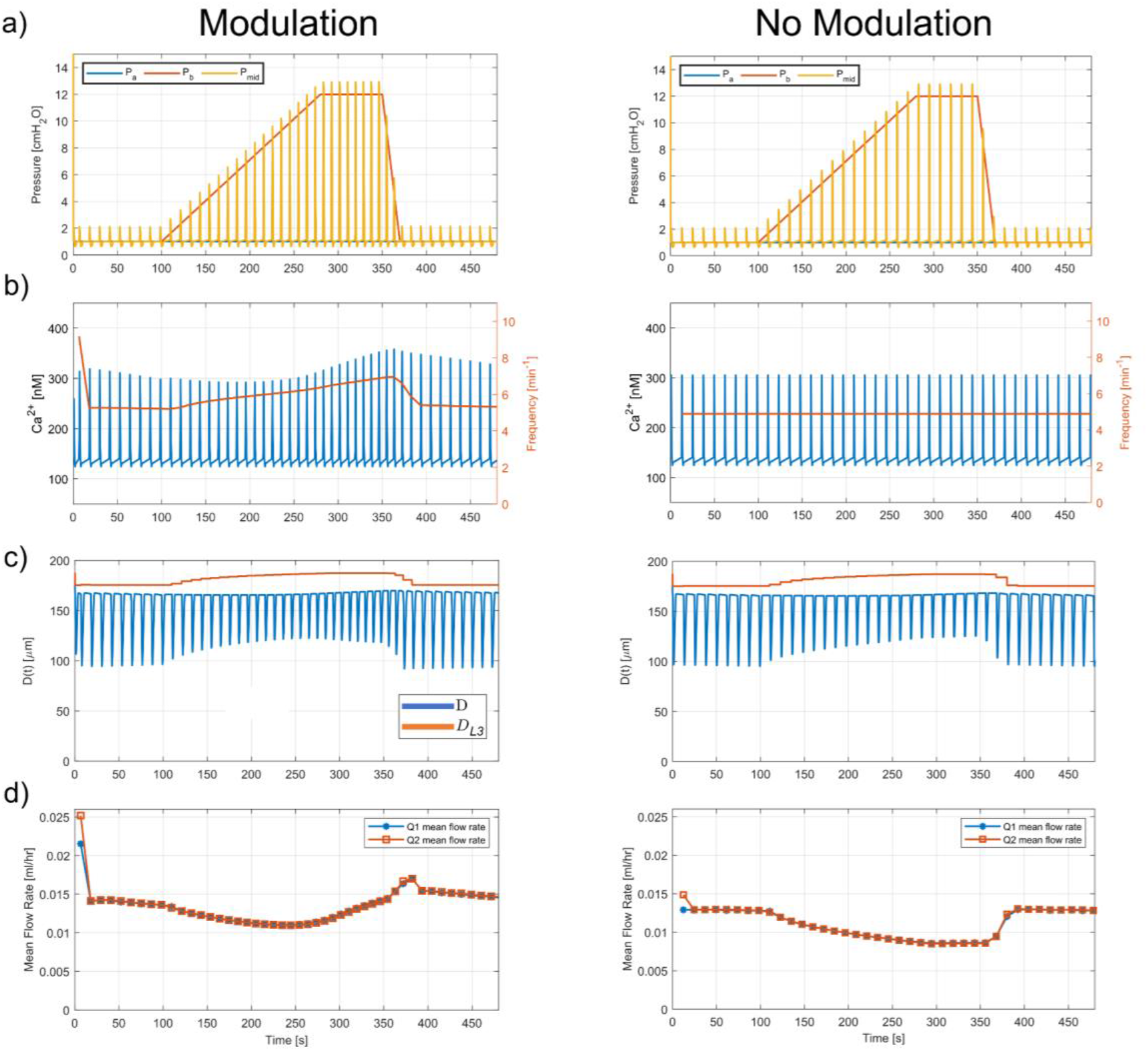
Plots of lymphangion parameters in the afterload pressure ramp simulation. **(a)** Inlet (*P*_*a*_) outlet (*P*_*b*_), and mid-lymphangion pressure (*P*_*mid*_), in afterload pressure ramp experiment. *P*_*mid*_ exceeds *P*_*b*_ during every systole. **(b)** Calcium concentration (blue) and contraction frequency (orange). Calcium concentration decreases initially, followed by an increase during the later stages of the pressure ramp. **(c)** Central lymphangion and passive downstream lymphangion (*D*_*L*3_) diameter in afterload pressure ramp experiment. Systolic central lymphangion diameter increases during the pressure ramp, due to increasing systolic mid-lymphangion pressure. Passive downstream diameter also increases steadily during the ramp. **(d)** Mean inlet and outlet flow rates.

A comparison with the model without the modulation algorithm is also shown (Figs. 7 and 8), where C_X_ and Ċ_d_ are fixed at the values corresponding to the TAWSS when Δ*P*_ab_ is set to zero, and Δ*P*_tm_ = 1cmH_2_O (Figs. 2a and 2b).

**Fig. 8.**
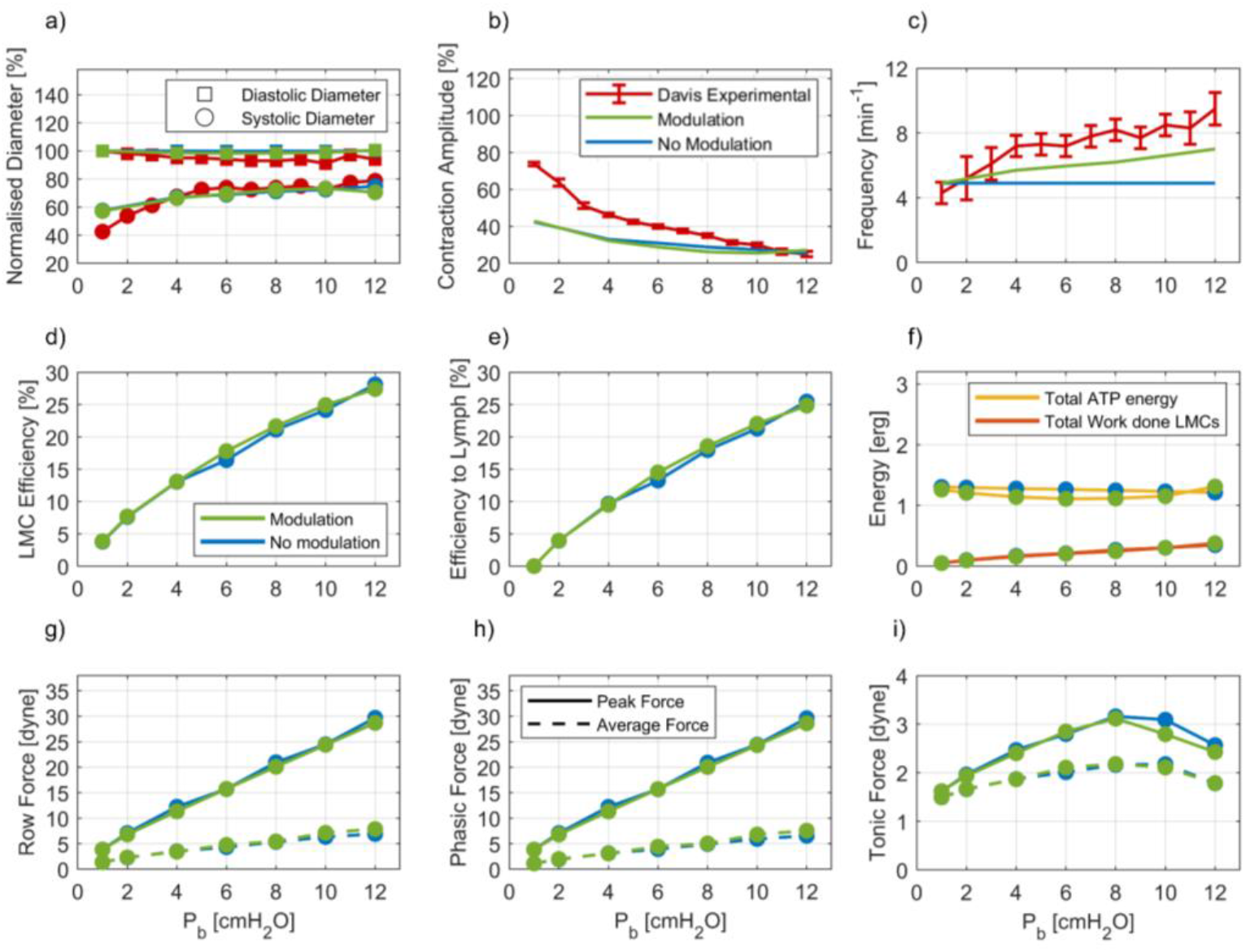
Analysis of parameters from a preload pressure ramp simulation, comparing modulated (green) and unmodulated (blue) with experiment (red). **(a)** Normalized EDD and ESD: model vs. experiment. **(b)** Contraction amplitude, model vs. experiment. **(c)** Contraction frequency, model vs experiment. **(d)** LMC efficiency in doing work (*η*_1_) as a function of outlet pressure *P*_*b*_. Efficiency increase is partially due to reduced outflow, hence there is less convection hence more myosin heads can complete the powerstroke. **(e)** Efficiency in producing lymph flow (*η*_2_). **(f)** Total ATP energy and total work done by LMCs as a function of outlet pressure *P*_*b*_. Total ATP energy follows the change in peak calcium concentration during the pressure ramp. Total work done by LMCs increases linearly as outlet pressure is increased. **(g)** Peak and average total phasic LMC force. **(h)** Peak and average cell force (force of a single LMC row). **(i)** Peak and average tonic LMC force.

**Fig. 9.**
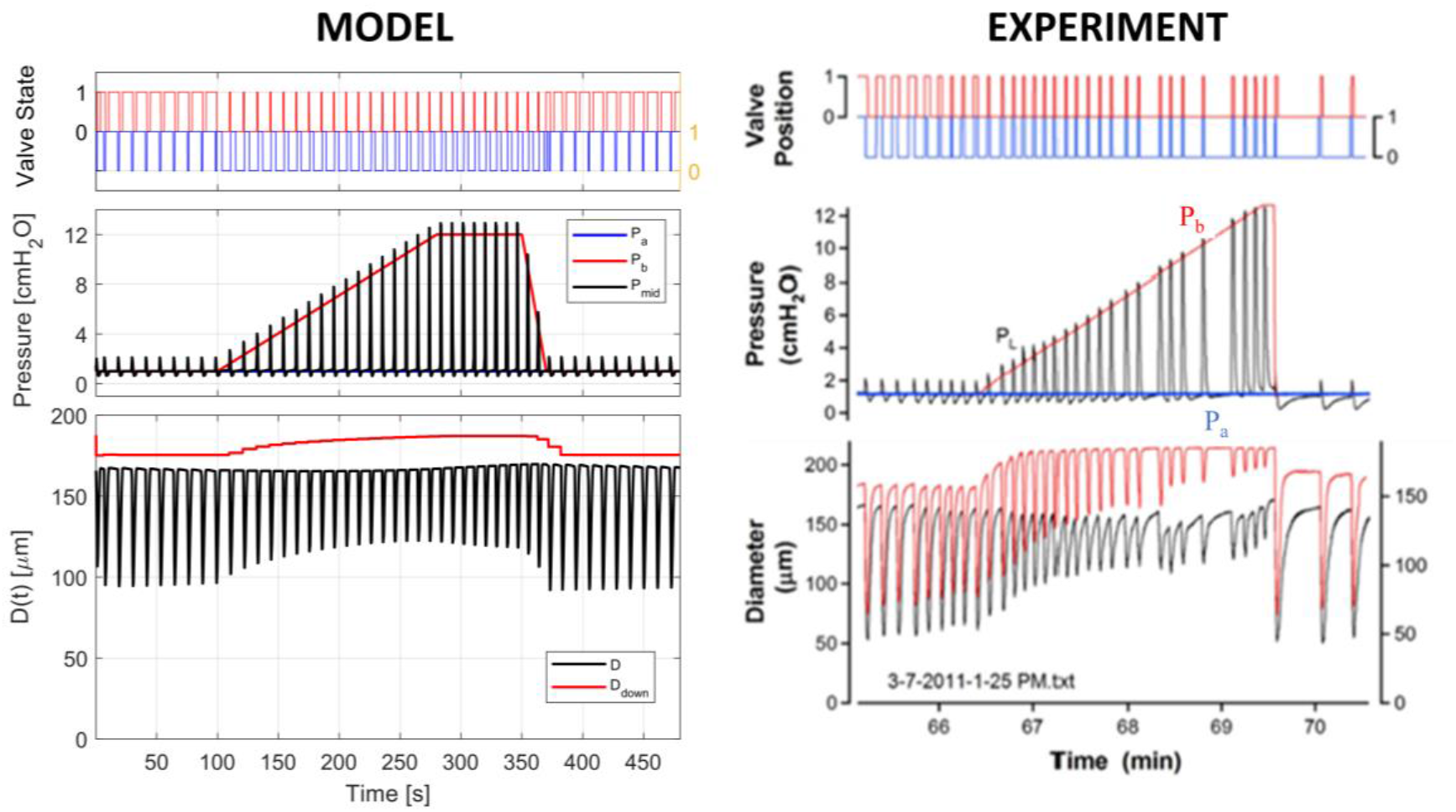
Comparison between afterload ramp simulation in model with modulation and an experimental afterload ramp [30]. Upper plots show valve state (1 – open, 0 – closed). Middle plots show the inlet, outlet, and mid-lymphangion pressures. Bottom plots show the central diameter and downstream lymphangion diameter (*D*_*L*3_). The downstream lymphangion is passive in the model simulations.

##### 2.8.2.2 Preload ramp simulation

In the actual inlet ramp experiment (section 3.4.2), because significant flow cause reduced contraction frequency and changes in contraction amplitude through NO production from the endothelium, two different sets of cannulating pipettes were used for protocols where inlet pressure P_a_ exceeded outlet pressure P_b_. High cannula resistances were preferred over the use of NO inhibitors, which could lead to irregular contraction patterns. To account for the high cannula resistance used in the experiments, the cannula resistance in the model was set to a value of 10^8^g/cm^4^s to minimize the effects of shear stress induced inhibition [38] (maximum cycle-average shear stress = 1.20cmH_2_O). The inlet pressure P_a_ was ramped from 2cmH_2_O to 16cmH_2_O at a rate of 3.5cmH_2_O/min. P_a_ was then held at 16cmH_2_O for 80s before decreasing back down to 2cmH_2_O. P_ext_ held at constant value of 0cmH_2_O, setting Δ*P*_tm_ = 2cmH_2_O when P_a_ = 2cmH_2_O. The model exhibits characteristic behaviour consistent with the experimental results (increasing contraction frequency and a decrease in contraction amplitude). Stretch signaling where r_Dd_ modulates the systolic peak calcium and r_Dd1_ modulates contraction frequency produced results that closely match the experimental results. Owing to the different rat mesenteric vessels used in experiments in the literature, each vessel naturally has a different passive pressure diameter relationship [32]. Therefore, we instead compare the end-diastolic (EDD) and end-systolic (ESD) diameters normalized to the diameter of the respective lymphangions at the initial conditions.

A comparison with the model without the modulation algorithm is also shown (Figs. 10 and 11), where C_X_ and Ċ_d_ are fixed at the values corresponding to the TAWSS when Δ*P*_ab_ is set to zero, and Δ*P*_tm_ = 2cmH_2_O (Figs. 2a and 2b).

**Fig. 10.**
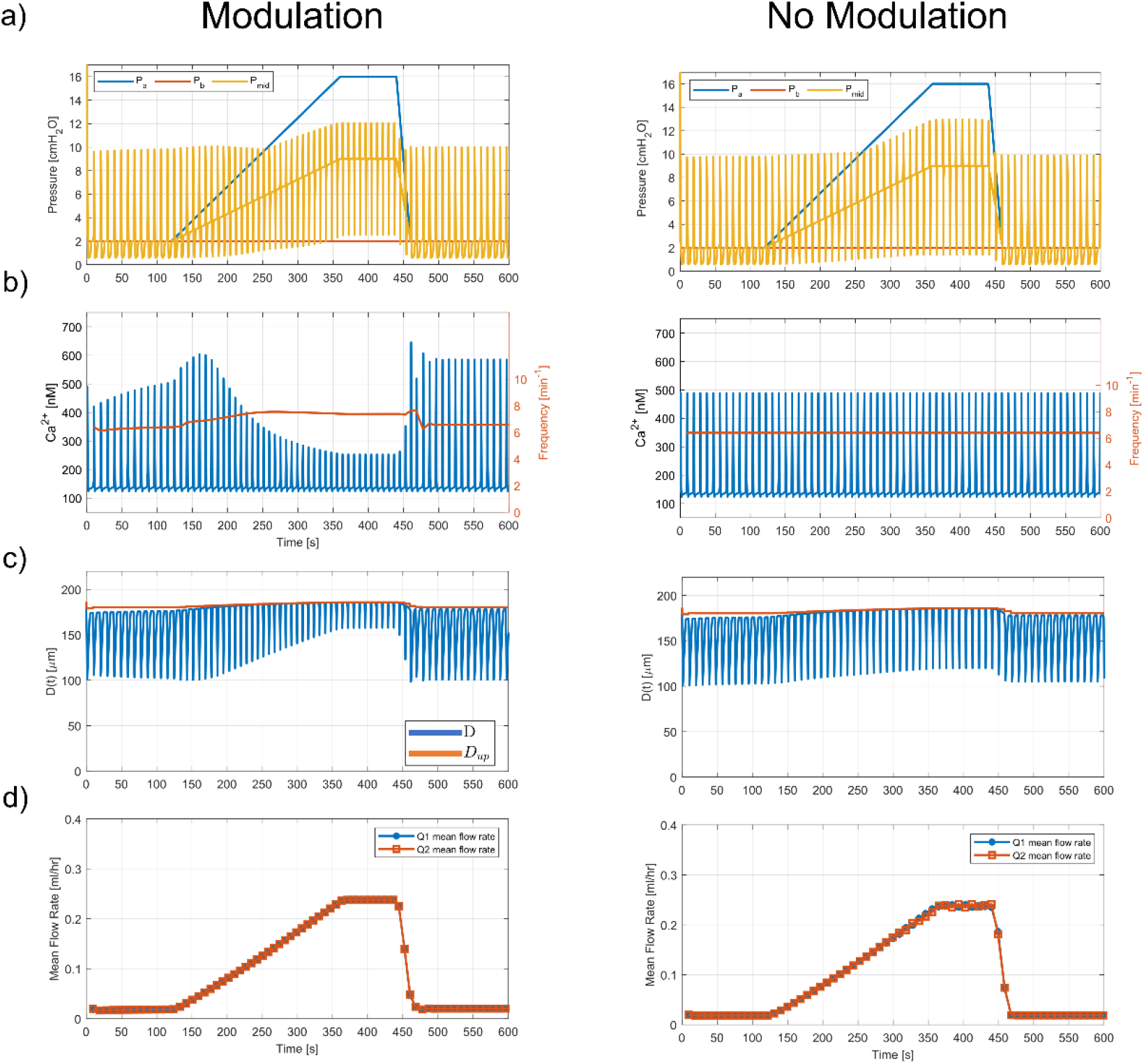
Plots of macroscopic parameters in the inlet pressure ramp simulation. **(a)** Inlet (*P*_*a*_), outlet (*P*_*b*_), and mid-lymphangion pressure (*P*_*mid*_). **(b)** Calcium concentration (blue) and contraction frequency (orange). Calcium concentration increases initially, but decreases during the later stages of the ramp. **(c)** Central lymphangion and passive upstream lymphangion diameter. **(d)** Mean inlet and outlet flow rates.

**Fig. 11.**
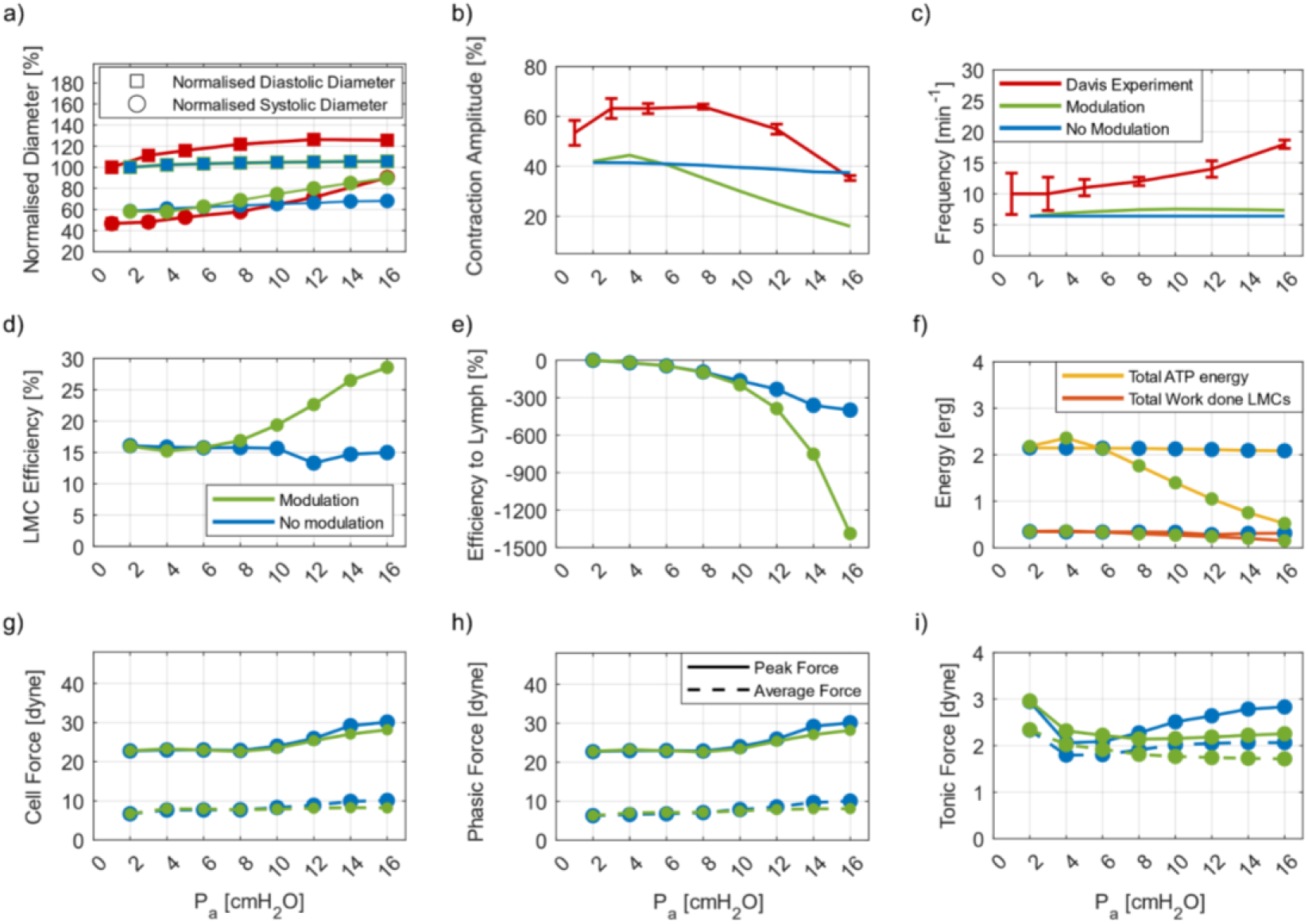
Analysis of parameters from a preload pressure ramp simulation, comparing modulated (green) and unmodulated (blue) with experiment (red). **(a)** Normalized EDD and ESD. **(b)** Contraction amplitude **(c)** Contraction frequency **(d)** LMC efficiency in doing work (*η*_1_) as a function of inlet pressure *P*_*a*_. Efficiency increase is partially due to decreased peak calcium, hence less ATP energy. **(e)** Efficiency in producing lymph flow (*η*_2_). **(f)** Total ATP energy and total work done by LMCs. Total ATP energy follows the change in peak calcium concentration during the pressure ramp. Total work done by the LMCs decreases as inlet pressure is increased. **(g)** Peak and average total phasic force of the LMC **(h)** Peak and average cell force (force of a single LMC row). **(i)** Peak and average tonic force of the LMC.

## 3 Results

### 3.1 Model verification with experimental data

#### 3.1.1 Lymphangion shear stress response

A key result in this set of simulations is the negligible difference in mean flow rate Q (Fig. 4l) between the modulated and unmodulated simulations but a decrease in energy produced from ATP hydrolysis and an increase in LMC efficiency (Fig. 4f).

Model parameters (eqs. 8–13) were first matched to experimental data [17,28] in the model verification stage. A comparison between the diameter traces of the mechanically modulated model lymphangion response and the experimental results of Gashev et al. showed strong positive correlation (a Spearman rank test between the systolic diameter values of model and results (*ρ*) = 1.00, p<0.05), and all five data points lie within the standard error of the mean (±SEM) from the data of Gashev et al. (Fig. 4a). The model response for Δ*P*_ab_ of 0cmH_2_O and 7cmH_2_O (Fig. 3a) shows a 28% increase in normalized systolic diameter and a 64% reduction in normalized contraction frequency. In the experimental results, there is a minor increase in the diastolic diameter with increasing Δ*P*_ab_ (Fig. 4a) (2% increase from 0 to 7 cmH_2_O), suggesting insignificant shear stress modulation of calcium during diastole. We have therefore based our shear modulation of tonic CEs only on systolic control parameters.

The mid-lymphangion pressure *P*_mid_ increased to a peak value of 6.2 cmH_2_O in the simulation with Δ*P*_ab_ = 0 cmH_2_O and 5.4 cmH_2_O in the simulation with Δ*P*_*ab*_ = 7 cmH_2_O (Fig. 3a). Minimum *P*_mid_, creating a suction effect for diastolic filling [2], was 1.5 cmH_2_O in the 7 cmH_2_O simulation and 2.5 cmH_2_O in the 7 cmH_2_O simulation. The duration of P_mid_ increase above the average pressure was longer in the 0 cmH_2_O case (1s vs 0.7s). The difference between the diameter and pressure traces can be attributed to the modulation of peak intracellular calcium concentration C_X_ and diastolic phase gradient Ċ_d_. C_X_ decreased by 52% (from 505nM to 242nM), resulting in a 28% increase in normalized systolic diameter (from 108μm to 150μm). Ċ_d_ is reduced from 2.55nM/s to 0.60nM/s, resulting in 64% reduction in normalized contraction frequency (from 8.8/min to 3.2/min) between the two simulations (Figs. 4b, 4d, and 4e). The flow rate (Fig. 3d) in the simulation with zero Δ*P*_ab_ shows outflow in systole, followed by inflow due to the suction effect. In the simulation with Δ*P*_ab_ = 7 cmH_2_O, systolic contractions increase the systolic outflow rate by a maximum of 0.4ml/hr above the mean diastolic flow rate.

In the case of zero Δ*P*_ab_, the phasic force increases to a maximum of 13.2 dyn in response to the increased calcium concentration, resulting in an almost identical peak cell force of 13.2dyn. The tonic force is maintained at a relatively constant value of 2.3 dyn during the contractile cycle (Fig. 3e). The tonic force remains relatively constant throughout the contractile cycle (maximum change of 25%) due to the tonic dashpot (Fig. 1D), resisting significant changes in tonic length. The tonic dashpot is thought to represent a combination of tonic contractile machinery and the fluid environment around the smooth muscle components. The phasic CE significantly contributes to the cell force during systole, whereas the tonic CE contributes significantly during the diastolic phase (Fig. 3e). As Δ*P*_ab_ is increased to 7 cmH_2_O, the model shows a decrease in peak phasic force to 11.7dyn, a peak cell force of 11.7dyn, and a decrease in average tonic force to 1.7dyn (Figs. 3e, 4g, 4h, and 4i). In the case of zero Δ*P*_ab_, LMCs converted approximately 8.8% of free energy from ATP liberation to work. This is compared to an efficiency of 13.0% with Δ*P*_ab_ = 7cmH_2_O (Fig. 4f). The efficiency increase is because of the reduced convection, allowing more myosin heads to bind to actin, so more heads can complete the powerstroke. The peak energy liberation rate was 3.3erg/s and the peak rate of muscle work was 0.42 erg/s for the simulation with Δ*P*_ab_ = 0cmH_2_O. A peak energy liberation rate of 1.0erg/s and a peak rate of muscle work of 0.29erg/s was found for the Δ*P*_ab_ = 7cmH_2_O (Fig. 4c). The decrease in peak energy rates is tied to the decrease in C_X_ because of modulation, which results in lower values of calcium saturations of troponin C and calmodulin, and hence lower phosphorylation and attachment rates [46]. The pump power (rate of energy transfer to moving lymph) is zero at zero Δ*P*_ab_, decreasing to an average of −1.7erg/s at Δ*P*_ab_ = 7cmH_2_O, as the favorable pressure gradient becomes a greater driving force to lymph flow. The rate of putting energy into moving lymph by the passive axial pressure difference is therefore +1.7erg/s.

Simulations in the absence of the modulation algorithm, where C_X_ and Ċ_d_ are fixed at constant values corresponding to no-flow conditions, and a transmural pressure of 5 cmH_2_O, show no significant changes in systolic diameter and contraction frequency (Figs. 4a and 4b). At Δ*P*_ab_ = 7cmH_2_O, a mean flow rate of 0.90ml/hr with modulation and 0.88ml/hr without modulation was observed, a 2% increase with modulation. This suggests no advantage for lymphangions to maintain strong active contractions during high flow situations (Figs. 4f and 4l). In the mechanically modulated case, the lymphangion relaxes to save metabolic energy; energy from ATP hydrolysis reduces from 2.15erg to 0.34erg for one contractile cycle between Δ*P*_ab_ = 0cmH_2_O to Δ*P*_ab_ = 7cmH_2_O (Fig. 4c). During high shear situations, the favorable pressure gradient forms a greater driving force to move lymph than the active lymph pump. The efficiency of LMCs remains relatively constant at 8.0% without modulation, whereas the efficiency increased to 13.0% at Δ*P*_ab_ = 7cmH_2_O with modulation (Fig. 4f). The peak cell and phasic forces are identical between the modulated and unmodulated results, suggesting that regulation of peak forces is independent of calcium (Figs. 4g and 4h). This is largely because, as the LMCs generate enough force to overcome the valve resistance and open the downstream valve, the LMCs begin to relax. The average tonic force is similar for the modulated and unmodulated lymphangion (within 5%) across all axial pressure differences.

#### 3.1.2 Lymphangion stretch response

A key result in this set of simulations suggests that there is an optimal transmural pressure Δ*P*_tm_ that induces maximum stretch-induced calcium mobilization. This is reflected in the plot of C_X_ (peak calcium concentration) modulation vs. r_Dd_ (diastolic stretch ratio), which shows a Gaussian curve behaviour (Fig. 2d). The peak LMC forces show a linear increase in the Δ*P*_tm_ range that we have tested. This behaviour is expected until pump failure, given that a greater cell force is required in higher pressure conditions to open the downstream valve and eject lymph. The highly nonlinear passive pressure-diameter behaviour [32] is likely an important factor in the reduction in contraction with increasing Δ*P*_tm_ above 3cmH_2_O, as this limits the strain of the lymphangion.

A comparison between the mechanically modulated model lymphangion response and the experimental results of Gashev et al. showed strong positive correlation (a Spearman rank test between the diastolic diameter values of model and results = 1.00, and systolic diameter correlation (*ρ*) = 0.90, p=0.08), and all diastolic and systolic normalized diameter points from simulation lie within standard deviations of the experimental data (Fig. 6a). In the experimental results, maximum contraction amplitude for rat mesenteric vessels was observed at Δ*P*_tm_ = 3cmH_2_O, followed by a reduction in contraction amplitude [28].

The mid-lymphangion pressure *P*_mid_ peaked at 4.1cmH_2_O in the simulation with Δ*P*_tm_ = 1 cmH_2_O and 6.2 cmH_2_O in the simulation with Δ*P*_tm_ = 3 cmH_2_O. *P*_mid_ then reduced to 2.6 cmH_2_O in the 1 cmH_2_O simulation and 2.9 cmH_2_O in the 3 cmH_2_O simulation (Fig. 5a) to create the suction effect. The suction amplitude increases from 0.4 cmH_2_O at Δ*P*_tm_ = 1 cmH_2_O to 2.0cmH_2_O at Δ*P*_tm_ = 3 cmH_2_O, and the duration of suction decreases from 2.4s to 0.8s. C_x_ increases from 300nM to 650nM when Δ*P*_tm_ increases from 1 to 3 cmH2O (Fig. 5c). C_x_ then decreases at higher Δ*P*_tm_, to 456nM at Δ*P*_tm_ = 3cmH_2_O (Fig. 6d). Ċ d increases from 1.02nM/s to 1.66nM/s, resulting in a 21% increase in normalized contraction frequency (from 4.8/min to 6.8/min) between the two simulations. Contraction frequency increases consistently with increasing levels of stretch (Fig. 6b). Flow rates (Fig. 5d) at Δ*P*_tm_ = 1 and 3 cmH_2_O show outflow in systole, followed by inflow due to the suction effect.

At Δ*P*_tm_ = 1 cmH_2_O, the phasic force increases to a maximum of 3.7 dyn in response to the increased calcium concentration, resulting in an almost identical peak cell force of 3.7 dyn. The tonic force is maintained at a relatively constant value, with an average of 1.6 dyn during the contractile cycle (Fig. 5e). Increasing Δ*P*_tm_ to 3cmH_2_O, the model shows an increase in peak phasic force to 8.54 dyn, a peak cell force of 8.6 dyn, and an increase in average tonic force to 2.2 dyn (Fig. 5e). In the 1cmH_2_O case, LMCs converted approximately 3.8% of free energy from ATP liberation to work. The efficiency increases to 5.5% when increasing Δ*P*_tm_ to 3cmH_2_O (Fig. 6f). The peak energy liberation rate was 3.1erg/s and the peak rate of muscle work was 0.13 erg/s for the 0cmH_2_O simulation. A peak energy liberation rate of 3.3erg/s and a peak rate of muscle work of 0.30 erg/s were found at Δ*P*_tm_ = 3cmH_2_O (Fig. 6c).

Simulations in the absence of stretch modulation, where C_X_ and Ċ_d_ are fixed at a constant value corresponding to no-flow conditions, and Δ*P*_tm_ of 1cmH_2_O, show an increase in both systolic and diastolic diameter with increasing Δ*P*_tm_. The normalized diastolic diameter increases from 81% to 90% (for both the modulated and the unmodulated model lymphangion), and the normalized systolic diameter increases from 48% to 57% (Fig. 6a). In the absence of stretch modulation, the stroke volume (EDV – ESV) decreases as Δ*P*_tm_ increases (from 42nL for 1cmH_2_O to 38nL at 5cmH_2_O) because of the larger force opposing the contractions of the LMCs. With the stretch modulation incorporated, as well as in experimental results of Gashev et al., maximum stroke volume is ejected at the Δ*P*_tm_ = 3cmH_2_O (58nL experimentally, 55nL in the model with modulation, and 39.8nL in the model without modulation). The efficiency of LMCs also increased with increasing levels of stretch, from 3.9% at Δ*P*_tm_ = 1cmH_2_O to 14.0% at Δ*P*_tm_ = 7cmH_2_O (Fig. 6f). The efficiency increases consistently with Δ*P*_tm_ because of a decrease in flow rate due to increased pressure load and hence a corresponding increase in the efficiency because of a decrease in myosin head convection (Fig. 6f). Efficiency for the modulated lymphangion is consistently smaller compared to the unmodulated lymphangion (in the modulated case, efficiency increased from 3.8% at 1cmH_2_O to 12.6% at Δ*P*_tm_ = 7cmH_2_O).

In the modulated case, the total ATP energy consumed in a contractile cycle increased from 1.34erg at Δ*P*_*tm*_ of 1cmH_2_O to 2.50erg at Δ*P*_*tm*_ = 3cmH_2_O, and then decreased to 1.98erg at Δ*P*_*tm*_ = 7cmH_2_O. The total work done by LMCs increased from 0.05erg at Δ*P*_*tm*_ = 1cmH_2_O to 0.25erg at Δ*P*_*tm*_ = 7cmH_2_O (Fig. 6c). The increase in work done by LMCs is due to the increasing pressure load and hence a higher resultant force generation. Energy liberation by ATP (Fig. 6c) follows the change in *C*_*X*_ modulation. The net increase in efficiency is therefore due to the fractional increase in the work done by LMCs being greater than the fractional increase in ATP energy liberated.

To investigate whether an optimal transmural pressure exists for the generation of active force, we plot the peak active force generated in a contractile cycle as a function of Δ*P*_*tm*_ for both the modulated and unmodulated model lymphangions (Fig. 6h). We find that there is a linear relationship between the peak active force generated by the LMCs and transmural pressure. A plot of row force (Fig. 5e) shows that there is an initial increase in force, where the systolic increase in calcium concentration leads to increased attachment of myosin heads to actin binding sites [46]. As the lymphangion starts to compress (decrease in diameter), the force generated by the LMCs eventually decreases as lymph is ejected from the downstream valve and the mid-lymphangion pressure *P*_*mid*_ decreases. The decrease in cell force allows the lymphangion to distend from systole (upwards slope in diameter) and allows the entry of lymph from the upstream lymphangion through the suction effect [2]. The fact that the peak phasic and peak cell forces are identical (maximum discrepancy 2.9% for Δ*p*_*ab*_ = 15cmH2O) between the modulated and unmodulated cases (Figs. 6g and 6h) suggests that the peak force is independent of peak calcium concentration, until pump failure. Once the LMCs generate enough force to overcome the valve resistance and open the downstream valve to eject lymph, the LMCs begin to relax.

The linear relationship between the maximum row force generated by the LMCs and Δ*P*_*tm*_ (Fig. 6g) suggests that the ability for myosin heads to form cross-bridges is not affected by increasing transmural pressure. The ability for myosin heads to generate contractile force during the isovolumetric contraction stage does not display the same Gaussian behavior as the calcium peak modulation, where peak calcium concentration increases to 3cmH_2_O and then drops off at higher Δ*P*_*tm*_ (Fig. 2d). This suggests that the decrease in peak calcium concentration, in combination with the increase in passive pressure due to strain stiffening, results in the fall in contractile amplitude and stroke volume that occurs from Δ*P*_*tm*_ = 3 to 7cmH_2_O [28].

### 3.2 Validation against pressure ramp experiments

#### 3.2.1 Afterload ramp experiment

Subjecting the model to a simulated gradual increase in outlet pressure P_b_ revealed the necessity of implementing inter-lymphangion stretch control. Because stroke volume is reduced when the outlet pressure is increased, the diastolic stretch ratio r_Dd_ decreases, which would normally cause a decrease in contraction frequency, as opposed to the positive chronotropic behaviour observed in experiments with afterload pressure ramps.

With implementation of inter-lymphangion stretch signaling, the calcium concentration profile shows a slight decrease in the peak calcium concentration during the P_b_ ramp, followed by an increase when P_b_ is larger than 8cmH_2_O, representative of the non-monotonic behaviour shown in (Fig. 2a). The peak calcium reached a minimum of 305nM and increased at the end of the ramp to 370nM. The contraction frequency also increases consistently during the ramp from 5/min to 7/min (Fig. 7b), a consequence of the diastolic stretch ratio based on the downstream lymphangion diameter (r_Dd3_) increasing from 2.16 to 2.25 as P_b_ rises from 1cmH_2_O to 12cmH_2_O.The intra-lymphangion stretch ratio r_Dd_ only increased from 2.10 to 2.13 over the same pressure range. The TAWSS 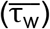 decreases from 0.45 to 0.23dyne/cm^2^ in the simulation as P_b_ is increased from 1cmH_2_O to 12cmH_2_O, leading to a 13% increase in contraction frequency (without considering effects due to stretch modulation) (Fig. 2b).

Comparing to the results without modulation, we observe that mean flow rate in the modulated case is consistently higher than the mean flow rate without modulation (Fig. 7d). Prior to the ramp, the mean flow rate is equal to 0.013ml/hr and decreases up to 0.011ml/hr. In the unmodulated case, the mean flow rate decreases from 0.013ml/hr prior to the ramp to 0.008ml/hr. This is in part due to the increase in contraction frequency with modulation (Fig. 7b) whereby the systolic flow rate becomes a larger component in the mean flow rate.

The end diastolic diameter (EDD) in the modulated case shows a paradoxical decrease in diameter, which may be a result of the muscle contracting more strongly when subjected to a slight increase in pressure (Fig. 7a). Supporting this theory, the mean tonic force (Fig/ 8i) increases from 1.5dyne to 2.2dyne, as P_b_ increases from 1 to 8cmH_2_O. This effect is independent of peak calcium concentration (C_X_), as the mean tonic force during diastole increases also in the unmodulated simulation (Fig. 8i), however, levels of calcium are similar in both simulations. We hypothesize that the increase in P_b_, by reducing outflow reduces CE convection, thereby facilitating more cross-bridge formation, and increasing the tonic force. The diameter of the central lymphangion displays a decreasing contraction amplitude during the outlet pressure ramp (Fig. 7c). As a result of these active (LMC contractions) and passive (passive lymph flow) phenomena, the normalized contraction amplitude decreased from 43% to 27% of initial diastolic diameter during the afterload ramp up to P_b_ = 12cmH_2_O. The contraction amplitude in the absence of mechanical feedback shows comparable values (Fig. 8b).

The experimental findings of Davis et al. [30] are visually similar to equivalent traces from the model results (Fig. 9). A comparison is made for the normalized EDD and ESD (Fig. 8a), normalized contraction amplitude (EDD - ESD) (Fig. 8b) and contraction frequency (Fig. 8c) comparing the model (green), the model without modulation (blue) and the experiment (red). We show the model with modulation has qualitatively the same trends as the experiment, notwithstanding that the model is matched to experimental data to different animals of the same species. The experiment shows a greater reduction in contraction amplitude during the pressure ramp, from 74% to 25% of the experimental diastolic diameter at *P*_b_ = 1 cmH_2_O, whereas in the model, contraction amplitude decreased from 43% to 27% of the simulated diastolic diameter at *P*_b_ = 1 cmH_2_O. The experimental contraction frequency increased from 4.3 to 9.5/min, whereas the model results show an increase from 5 to 7/min.

The efficiency of the LMCs increases with increasing outlet pressure P_b_, reaching a peak value of 27% at P_b_= 12cmH_2_O, compared to 4% at the beginning of the pressure ramp when P_b_ = 1cmH_2_O (Fig. 8d). The efficiency of lymph propulsion (calculated using the pump power equation) increases similarly from 0% when P_b_ = 1cmH_2_O to 24.4% when P_b_ = 12cmH_2_O. The total work done increases linearly with P_b_ as the force required to overcome the adverse pressure difference increases (Figs. 8g and 8h). The increase in force demand is shown by the increase in peak phasic and row forces as P_b_ is increased (Fig. 8g and 8h). Peak phasic force increases from 3.6dyn to 27.2dyn across the pressure ramp. Peak total cell force follows peak phasic force, also increasing from 3.6dyn to 27.2dyn. The tonic force behaves in the opposite manner to the total ATP energy (Fig. 8f and 8i). The increase in average tonic force is a result of the increase in diastolic pressure, as well as reduced convection from the decreased flow rate (Fig. 7d), allowing more myosin heads to bind to actin, so more heads can complete the powerstroke facilitating the formation of more tonic cross-bridges (with help from the effects of the tonic dashpot). The flow rate profile during the P_b_ ramp (Fig. 7d) shows the opposite response to the change in average tonic force, supporting this explanation.

When P_b_ is held at the highest pressure of 12cmH_2_O, the peak calcium concentration increases steadily, approaching a constant value. Due to the nature of the modulation algorithm, whereby the previous contractile cycle is used as the variable to control the lymphangion behaviour in the current cycle, there is a lag of a few contractile cycles to reach a new steady state. This period of stabilization could physiologically represent the time required for the lymphangion to adjust into a consistent response upon a change in physical conditions. In published observations, constant pressure conditions are typically held for a minimum of 10 contractions for the vessel to stabilize its response [30].

Overall, the afterload ramp simulation exhibits minor changes in wall shear stress and insignificant changes in intra lymphangion stretch during diastole. The implementation of stretch modulation with r_Dd_ modulating C_X_ and inter-lymphangion control r_Dd3_ to modulate Ċ_d_ suggests that modulation increases the contraction frequency during the pressure ramp.

#### 3.2.2 Preload ramp simulation

A preload ramp experiment tests the response of the lymphangion to a favorable pressure difference by exposing single, isolated lymphangions to selective changes in preload [38]. As in the afterload ramp simulation, by considering only the stretch signaling from the central lymphangion, the slight decrease in contraction frequency (mainly due to inhibition due to shear stress) during the inlet pressure ramp simulation is unrepresentative of the positive chronotropic behaviour observed in experiments.

With implementation of interlymphangion stretch signaling, the calcium concentration profile shows an initial increase in the peak calcium concentration during the P_a_ ramp, followed by a decrease as P_a_ rises above 4cmH_2_O. The peak calcium reaches a maximum of 630nM when P_a_ = 4cmH_2_O and decreases at the end of the ramp to 260nM due to both the increase in shear stress which decreases C_X_ (Fig. 2a), and the increase in r_Dd1_ also decreasing C_X_ as the stretch ratio reaches the other side of the Gaussian modulation curve (Fig. 2b). The stretch ratio involving the upstream passive lymphangion r_Dd1_ increases from 2.23 to 2.35 as P_a_ is increased from 2cmH_2_O to 16cmH_2_O (Fig. 10a). The local stretch ratio r_Dd_ shows a similar increase from 2.21 to 2.33 over the same pressure range, as the upstream valve is open in diastole during the preload ramp. The TAWSS 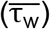 increases from 0.50 to 1.2dyn/cm^2^ in the simulation as P_a_ is increased from 2 to 16cmH_2_O. The contraction frequency increases consistently during the preload ramp, from 6.2 to 7.6/min (Fig. 11c), due to the increase in diastolic stretch ratio r_Dd1_, being greater than the inhibitory effect of the increase in 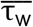. The preload pressure ramp therefore tests the lymphangion response to both shear stress and stretch.

Comparing to the results without modulation, we observe that mean flow rate in the modulated case is consistently within 1% of the mean flow rate without modulation (Fig. 10d). Prior to the ramp, the mean flow rate is equal to 0.018ml/hr, and it increases to a maximum of 0.238ml/hr. In the unmodulated case, the mean flow rate increases from 0.018ml/hr prior to the ramp up to 0.239ml/hr (Fig. 10d). The maximum flow rate therefore is largely independent of modulation (<1% change), from which we can draw a similar conclusion to that drawn from the simulations of varying axial pressure difference (Δ*P*_*ab*_), that passive flow becomes a much larger driving force in lymph flow than active contractions when the pressure gradient is favourable.

Comparisons are made in Fig. 11 for the normalized EDD and ESD (Fig. 11a), normalized contraction amplitude (Fig. 11b) and contraction frequency (Fig. 11c) between the model (green), the model without modulation (blue) and experiment (red). The diameter trace of the central lymphangion displays a decreasing contraction amplitude during the inlet pressure ramp, resulting in increasing ESD (Fig. 11a). As a result of the active and passive effects, the contraction amplitude decreased during the preload ramp from 73μm at P_a_ = 2cmH_2_O to 28μm at P_a_ = 16cmH_2_O (Fig. 10c). In the absence of modulation, the contraction amplitude displays a smaller decrease, from 73μm at P_a_ = 2cmH_2_O to 66μm at P_a_ = 16cmH_2_O (Fig. 10c). The experiment shows a reduction in normalized contraction amplitude from 53% to 35% of the initial diastolic diameter. The experimental contraction frequency increased from 10 to 18/min whereas the model results show an increase from 6.2 to 7.6/min. In the absence of 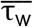 modulation (which acts to decrease contraction frequency), the model contraction frequency increases from 6.2 to 12/min over the same pressure range, showing that the shear modulation largely inhibits the increase in contraction frequency.

Comparison is made (Fig. 12) between the responses of an experimental lymphangion and of the model to a preload ramp. They show similar characteristic responses in the central lymphangion. This includes an increase in ESD and an increase in contraction frequency during the afterload pressure ramp. The mid-lymphangion pressure trace reveals that the peaks during systolic contraction and the suction pressure amplitude is much larger in the model compared to the experimental data.

**Fig. 12.**
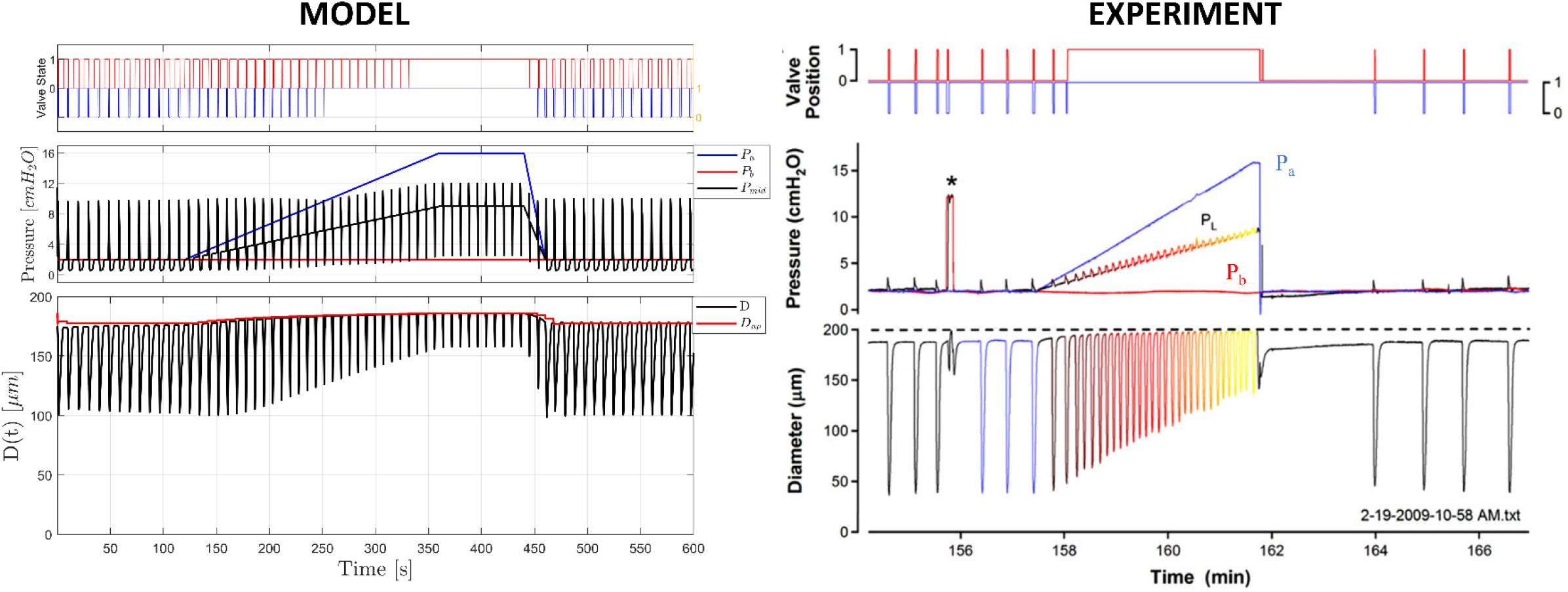
Comparison between preload simulation performed with model (left) and an experimental preload ramp [38]. Upper plots show valve state (1 – open, 0 - closed) for inlet (blue) and outlet (red) valves. Middle plots show inlet (blue), outlet (red), and mid-lymphangion (black for model, varying for experiment) pressures. Left lower plot shows model central diameter (black) and passive downstream diameter (red). Color scheme in the experimental diameter trace (right lower plot) indicates the variation in mid-lymphangion pressure from low (blue) to high (yellow).

The efficiency of the LMCs increases with increasing inlet pressure P_a_, rising from 15% when P_a_ = 2cmH_2_O to 29% when P_a_ = 16cmH_2_O. Without modulation, the efficiency decreases from 15% to 13% over the same pressure range (Fig. 11d). This is due to the maintained C_X_ levels in the preload ramp, producing diminishing returns in the work done by LMCs. Maintaining active contractions in situations of high favorable axial pressure is therefore unnecessary and provides insignificant change in flow rate (Fig. 10d). The total ATP energy consumed over a contractile cycle follows the modulation of peak calcium C_X_, where there is an increase up to P_a_ = 4cmH_2_O, followed by a consistent decrease for the modulated case. In the absence of modulation, the total ATP energy is fixed at 2.1erg (Fig. 11f). The total work done decreases with increasing P_a_, as the force required by the lymphangion is decreased due to shear inhibition, and passive flow becomes a greater facilitator to lymph flow (Fig. 11f). The peak cell force remains relatively constant at 23dyn as P_a_ is increased up 10cmH_2_O (Figs. 11g and 11h). The peak cell force then increases to 28dyn when P_a_ reaches 16cmH_2_O. The peak phasic force (total phasic force produced by LMC) follows the increase in peak cell force. The mean tonic force decreases throughout the pressure ramp, from 2.35dyn to 1.71dyn (Fig. 11i).

Scallan et al. [38] suggested that key contraction parameters during inlet pressure ramps were “very similar in pattern and magnitude to those at 4.5-5min in response to inlet pressure steps”. The key parameters of normalized contraction amplitude and contraction frequency are compared between the model and the experiment in Fig. 11. Normalized diameter is used to account for the differences in passive pressure-diameter relationships of the lymphatic vessels. Experimentally there is a decrease in contraction amplitude (Fig. 11b) with increasing P_a_, and increasing contraction frequency (Fig. 11c). Our model shows qualitatively the same behaviour, considering that the model was matched to results from a different animal [17,28]. The qualitative agreement between the model simulations and the experimental results is therefore notable.

## 4 Discussion

In this study, we have incorporated the role of mechanical feedback into a computational model capturing subcellular mechanisms of lymphatic muscle contraction [46], coupled with a well-characterized macroscale lumped-parameter model, producing pressure, diameter and flow parameters representative of experimental results performed on rat mesenteric lymphatics in vitro. Using experiments performed by Gashev et al. [17,28] as a basis for parameter values for the regulation of calcium amplitude and frequency in the model, we can investigate how intracellular calcium changes in response to shear stress and stretch. The model benefits from simplicity, directly relating physical conditions experienced by the lymphangion to quantitative changes in intracellular calcium. This model is the first to study changes in intracellular calcium in lymphatics in response to mechanical load (experimental or computational) and provides insight into the subcellular contractile machinery (phasic and tonic CE forces) and LMC efficiency, and how they are affected by mechanical signals. We do not directly model the cellular mechano-transduction pathways or the voltage-dependent calcium channels and their respective ion fluxes, as much of the identity and importance of different ion channels in response to mechanical signals is not well characterized [19]. We hope the model can provide insight into the qualitative and quantitative effects of mechanical signals on intracellular calcium and inspire further experimental investigations in the future.

The pathway of shear-stress-induced calcium regulation is well recognized as being mediated by NO and histamine produced by lymphatic endothelial cells, acting through pathways that limit Ca^2+^ influx. However, the endothelial mechanobiology and the transduction mechanism of shear stress signals in lymphatics is not well characterized. In blood-vessel endothelial cells, structures such as the glycocalyx component and adhesion molecules are known to be transducers of shear stress signals [55]. A better understanding of these structures and pathways in lymphatics can help better characterize the effect of various EDFs on lymphatic contractions. Currently, the model lumps the effect of shear-stress-derived EDFs as a normalized value from 0-1, bypassing the modeling of the exact concentrations of NO produced, due to a lack of experimental data as well as a lack of quantitative understanding of the interactions between NO and [Ca^2+^]. Experiments measuring NO have been performed in vascular endothelial cells [56]. Additional experiments measuring NO in response to shear stress, as well as a better understanding of the interaction between NO and calcium, would be beneficial to further developments in sub-cellular models of lymphatics in the future.

Limited experimental studies have been done on the relationship between stretch-induced calcium mobilization and lymphatic pump activity and there are conflicting data between different studies [12,33,34]. Shirasawa et al. [33] found in rat thoracic duct that incremental stretch increased the frequency of phasic contractions, as well as the amplitude of the isometric force, but not the basal level or amplitude of the calcium transient. Additional experiments on calcium fluorescence imaging will help provide quantitative measurements of how stretch regulates calcium and are important for understanding stretch regulation in lymphatics. Stretch-activated ion channels have been explored in response to mechanical stretch in endothelial cells [57]. The exact identity of the stretch-activated calcium channels in LMCs is unknown, but Piezo1 is a candidate protein that transduces mechanical forces into cationic currents in blood vessels. The model makes a prediction of the calcium amplitude response to stretch as following a Gaussian behaviour (Figs. 2b and 2e), however this is not well testified by experimental data [12,33,34]. Experimental studies exploring the exact identity of the stretch-activated calcium channel could potentially help explain the behaviour of the calcium regulation curves in response to both stretch and shear (Figs. 2a, 2c and 2d).

The model was tested using pressure ramp experiments performed by Davis, Scallan and co-workers [30,38] to test its response to adapting pressure conditions, where a combination of shear stress and stretch modulation is involved. The model response, with mechanical regulation matched to experimental data from Gashev et al. shows a weaker contractile ability compared to those tested by Davis, Scallan, and co-workers, potentially because they used a single-lymphangion preparation with two valves, in comparison to the 0.8-1.0 cm segments without valves that Gashev et al. used. Given the variations between circumstances such as the lymphatic vessel, there will be variations in the responses between experiments in different studies. The degree of reproducibility obtained between the model and the experimental results is therefore exceptional, allowing us to probe insights into the microscopic mechanisms occurring in the lymphatic cell, as well as parameters such as pressure, flow rate, LMC efficiency etc. In the afterload experiment, we observed a net increase in efficiency, due to the increased work done by the LMCs in contracting to overcome the adverse pressure difference. The peak calcium C_X_ remained relatively constant through the afterload pressure ramp, as the diastolic stretch ratio remains relatively constant, and the contraction frequency increased, due to the incorporation of the downstream diameter signaling in frequency modulation. Davis et al. [30] hypothesized that a decrease in EDD as a function of *P*_b_ is analogous to the underlying increase in contractility associated with myogenic constriction in arterioles. We confirm this hypothesis by showing an increase in mean tonic force, due to the reduced CE convection with increasing downstream pressure, facilitating more cross-bridge formation, subsequently increasing the smooth muscle (tonic) force. In the preload experiment, we observed an initial increase followed by a decrease in peak calcium C_X_, where the initial increase is due to the stretch, and the following decrease is due to shear-stress-induced inhibition and stretch reaching levels corresponding to the right side of the Gaussian control curve (Fig. 2d). We observe a net increase in LMC efficiency from 15% to 29%, as the fractional decrease in total ATP energy is greater than the decrease in work done by the LMCs. In comparison, in the simulation without modulation, the efficiency decreases from 15% to 13% during the preload ramp. The model incorporated passive upstream and downstream lymphangions in these pressure ramp simulations to explore the potential effects of signaling between lymphangions in the ramp experiments. We found that close replication to the experimental results was obtained when stretch in the downstream lymphangion (in the afterload experiment) and upstream lymphangion (in the preload experiment) regulates the contraction frequency of the central lymphangion. This reveals the importance of signaling between lymphangions, as reported in the literature [44,53].

Lymphatic vessels perform both rapid, rhythmical phasic contractions as a primary mechanism to generate flow and slow tonic constrictions by lymphatic smooth muscle in order to regulate flow via diameter-based resistance. This study shows interesting findings of how these phasic and tonic forces change as shear and stretch are modulated. With shear modulation incorporated, we observe a decrease in mean phasic and cell force due to a reduction in C_X_. With stretch modulation incorporated, we observe no significant change in peak and mean cell and phasic forces. This is attributed to the relaxation of the LMCs after subsequent opening of the downstream valves, expelling lymph and decreasing P_mid_. The increased C_X_ due to stretch modulation contributes to a greater peak and average tonic force. Quantitative estimates of the force contributions from the phasic and tonic CEs in the model and how they are affected by mechanical regulation further our understanding of their contributions during different stages of the contractile cycle. It has been shown experimentally that shear-reduced tone and stronger phasic contractions result in more energy-efficient pumping [58]. A better understanding of LMC contractile properties will help us devise potential treatments to promote flow in lymphedema.

LMC efficiency was first explored computationally by Morris et al. [46] under conditions where the effect of mechanical regulation was not incorporated. The present study makes further estimates of LMC efficiency under various mechanical conditions, as well as testing the effect of incorporating mechanical regulation. We observe that efficiency increases with shear regulation and decreases due to stretch modulation (Figs. 4f and 6f). LMC efficiency increases to 28 and 30% in the afterload and preload experiments. However, there are no experimental comparisons that we can make with LMCs. Experimentally, ATP usage is measured by monitoring the concentration of inorganic phosphate via fluorescent protein MDCC_PBP, or via the amount of heat generation. Thermodynamic efficiency has been reported to be around 20% for cardiac muscle [59], and 18% for vascular smooth muscle [60]. The ability to culture LMCs and measure efficiency would be valuable in validating the values obtained in the model.

A lack of experimental data directly measuring calcium in response to mechanical signals prevents us to ensure quantitative accuracy of intracellular calcium concentration in the LMCs. The calcium concentrations on which we based the model were obtained using the fluorescence ratio between two different intensities [12]. It has been shown that the fluorescence ratio is not a precise measurement for [Ca^2+^] [61]. However, as the values of calcium concentration and the predicted calcium saturation values we used in the model produce the macroscopic response shown in experiments, we are confident that the relative changes in calcium concentration are representative of biology.

There are also potential inaccuracies in the model shear stress calculations calculated from the data. Gashev et al. [17] used mesenteric lymphatic segments (0.8 to 1.0cm) without valves cannulated with resistive pipettes in order to impose an axial pressure difference across the segment. The passive pressure-diameter relationship of the lymphangions used in the experiment was also different to that assumed in the model. As the paper did not report a specific resistance (or diameter) of the pipettes, we therefore used values quoted by Scallan et al. [38] in our calculations. For the purposes of modeling and understanding the effect of mechanics on lymphatic response, the exact value of shear stress experienced is of lesser importance. Due to biological variability, shear stress may elicit different responses in different lymphatics. The more essential objective is that the model captures the linear relationship between shear stress and axial pressure difference (due to the low Reynolds number of the flow) and produces a macroscopic response that matches the experimental data of Gashev et al. [17]. The ranges of shear stress calculated in the model are comparable to values of shear stress previously measured in the literature [62]. More experiments directly monitoring the shear stress on lymphatic endothelial cells during flow conditions can allow a more accurate estimation of the true shear stress experienced by cells in the lymphangion.

More experimental data on lymph pumping experiments performed at a greater range of physical conditions exposing the lymphangion to different combinations of stretch and shear signaling can improve the 3D control algorithms we have constructed for the model. The surface plots (Figs. 2a and 2b) show the extrapolated curve fits in relation to the available experimental data [17,28]. As a result, the model can only be deemed accurate within the proximate ranges of the experimental data. The mathematical framework set by the model can still be used, but more experimental data would help further improve the accuracy of the model.

## 5 Conclusion

The model consists of the implementation of a single central lymphangion surrounded by passive non-contracting upstream and downstream lymphangions. Extending the scale of the model to multiple lymphangions (chain) and networks can allow for a more realistic model of the lymphatic network. This involves computational complexity due to multi-scale coupling; however, a few suggestions are outlined to resolve this issue [46]. Parameter sensitivity analysis can also be conducted on this new sub-cellular model of lymphatics [51]. Interesting behaviors, and a greater understanding of the effect of lymphatic parameters relating to the subcellular machinery could pave the way for identifying new interventions to target specific components of lymphatic cells.

In conclusion, this study has integrated mechanical feedback mechanisms into a sub-cellular model of the lymphangion by coupling mechanical signals experienced by the lymphangion to quantitative changes in intracellular free calcium ion concentration. We started by verifying the model framework by matching axial pressure difference (Δ*P*_*ab*_) and transmural pressure (Δ*P*_*tm*_) experiments performed [17,28]. The model was then validated with separate pressure ramp experiments to explore the dynamic effects of shear and stretch signaling. The model can be used to investigate parameters that are difficult to measure experimentally, including flow rate, pressure, forces in subcellular LMC components, phasic and tonic CEs, and LMC efficiency. Understanding the molecular functioning of LMCs is necessary for greater understanding of the system’s performance [63] and is useful for identifying potential pharmaceutical interventions. The ability to culture LMCs while maintaining their contractile response is important in advancing our understanding of their contractile properties and how they respond to physical stimuli.

## Acknowledgements

The authors acknowledge Drs Daniel Watson and Jennifer Frattolin for their valuable advice and guidance on the manuscript. This research study was funded by NIH grant U01-HL123420 and the Sir Leon Bagrit Trust. CDB acknowledges funding from NIH grant R01-HL-122578.

## Appendix

**Table 1:**
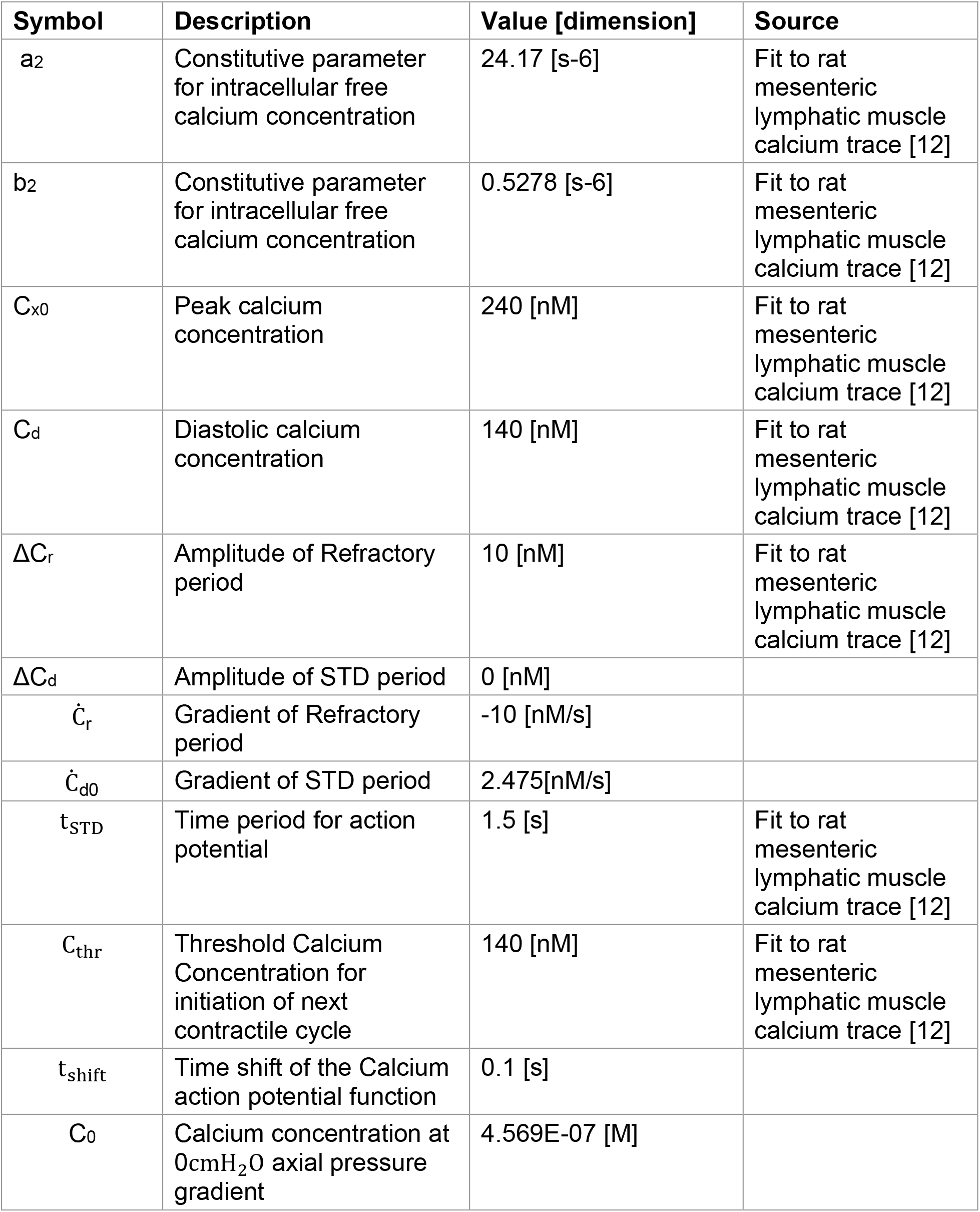
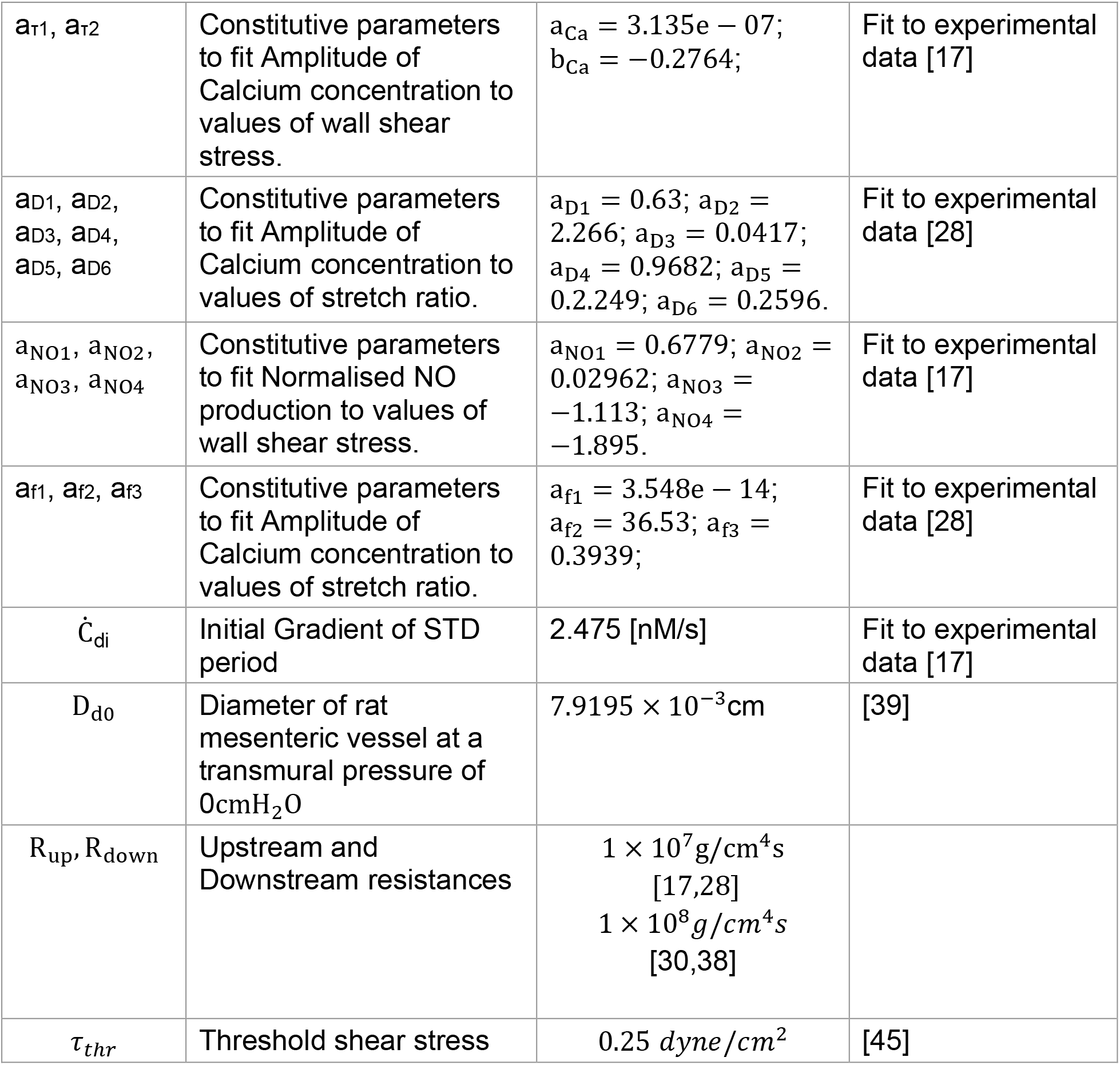
Parameter Values.

| Symbol | Description | Value [dimension] | Source |
| --- | --- | --- | --- |
| $a_2$ | Constitutive parameter for intracellular free calcium concentration | 24.17 [s-6] | Fit to rat mesenteric lymphatic muscle calcium trace [12] |
| $b_2$ | Constitutive parameter for intracellular free calcium concentration | 0.5278 [s-6] | Fit to rat mesenteric lymphatic muscle calcium trace [12] |
| $C_{x0}$ | Peak calcium concentration | 240 [nM] | Fit to rat mesenteric lymphatic muscle calcium trace [12] |
| $C_d$ | Diastolic calcium concentration | 140 [nM] | Fit to rat mesenteric lymphatic muscle calcium trace [12] |
| $\Delta C_r$ | Amplitude of Refractory period | 10 [nM] | Fit to rat mesenteric lymphatic muscle calcium trace [12] |
| $\Delta C_d$ | Amplitude of STD period | 0 [nM] | |
| $\dot{C}_r$ | Gradient of Refractory period | -10 [nM/s] | |
| $\dot{C}_{d0}$ | Gradient of STD period | 2.475[nM/s] | |
| $t_{STD}$ | Time period for action potential | 1.5 [s] | Fit to rat mesenteric lymphatic muscle calcium trace [12] |
| $C_{thr}$ | Threshold Calcium Concentration for initiation of next contractile cycle | 140 [nM] | Fit to rat mesenteric lymphatic muscle calcium trace [12] |
| $t_{shift}$ | Time shift of the Calcium action potential function | 0.1 [s] | |
| $C_0$ | Calcium concentration at 0cmH<sub>2</sub>O axial pressure gradient | 4.569E-07 [M] | |
| $a_{\tau 1}, a_{\tau 2}$ | Constitutive parameters to fit Amplitude of Calcium concentration to values of wall shear stress. | $a_{Ca} = 3.135e - 07;$<br>$b_{Ca} = -0.2764;$ | Fit to experimental data [17] |
| $a_{D1}, a_{D2},$<br>$a_{D3}, a_{D4},$<br>$a_{D5}, a_{D6}$ | Constitutive parameters to fit Amplitude of Calcium concentration to values of stretch ratio. | $a_{D1} = 0.63; a_{D2} =$<br>$2.266; a_{D3} = 0.0417;$<br>$a_{D4} = 0.9682; a_{D5} =$<br>$0.2.249; a_{D6} = 0.2596.$ | Fit to experimental data [28] |
| $a_{NO1}, a_{NO2},$<br>$a_{NO3}, a_{NO4}$ | Constitutive parameters to fit Normalised NO production to values of wall shear stress. | $a_{NO1} = 0.6779; a_{NO2} =$<br>$0.02962; a_{NO3} =$<br>$-1.113; a_{NO4} =$<br>$-1.895.$ | Fit to experimental data [17] |
| $a_{f1}, a_{f2}, a_{f3}$ | Constitutive parameters to fit Amplitude of Calcium concentration to values of stretch ratio. | $a_{f1} = 3.548e - 14;$<br>$a_{f2} = 36.53; a_{f3} =$<br>$0.3939;$ | Fit to experimental data [28] |
| $\dot{C}_{di}$ | Initial Gradient of STD period | 2.475 [nM/s] | Fit to experimental data [17] |
| $D_{d0}$ | Diameter of rat mesenteric vessel at a transmural pressure of 0cmH<sub>2</sub>O | $7.9195 \times 10^{-3} \text{cm}$ | [39] |
| $R_{up}, R_{down}$ | Upstream and Downstream resistances | $1 \times 10^7 \text{g/cm}^4 \text{s}$<br>[17,28]<br>$1 \times 10^8 \text{g/cm}^4 \text{s}$<br>[30,38] | |
| $\tau_{thr}$ | Threshold shear stress | 0.25 *dyne/cm*<sup>2</sup> | [45] |

## Footnotes

1 In the model, the trigger of contractions is the intracellular calcium concentration. In a muscle cell, the periodic sudden massive influx of calcium into the cytosol via L-type channels is what causes the action potential. Dividing the calcium waveform into three phases (named after analogous phases in cardiac contraction), we here treat it analogously to the conventional physiological description of the electrical events leading to and including the action potential. Modelling of the coupled electrical and calcium flux events involved in LMC pacemaker oscillations has been conducted (Hancock et al. [64]). Although it would be ideal to couple such an autonomous oscillator into the model, we do not do so here.

2 For low-Reynolds-number flow, wall shear stress is linearly proportional to the axial pressure difference Δ*P*_*ab*_. The constant of proportionality between the two variables is linearly dependent on the vessel diameter and length. The set of isolated lymphatic vessels tested in Gashev et al. [17,28] reported the mean response. In these simulations, we use values of shear stress for the model lymphangion. It should be noted that shear-inhibition behaviors may differ between lymphatics of different sizes.

